# Reliability and sensitivity of two whole-brain segmentation approaches included in FreeSurfer – ASEG and SAMSEG

**DOI:** 10.1101/2020.10.13.335737

**Authors:** Donatas Sederevičius, Didac Vidal-Piñeiro, Øystein Sørensen, Koen van Leemput, Juan Eugenio Iglesias, Adrian V. Dalca, Douglas N. Greve, Bruce Fischl, Atle Bjørnerud, Kristine B. Walhovd, Anders M. Fjell, for the Alzheimers Disease Neuroimaging Initiative

## Abstract

An accurate and reliable whole-brain segmentation is a key aspect of longitudinal neuroimaging studies. The ability to measure structural changes reliably is fundamental to detect confidently biological effects, especially when these affects are small. In this work, we undertake a thorough comparative analysis of reliability, bias, sensitivity to detect longitudinal change and diagnostic sensitivity to Alzheimer’s disease of two subcortical segmentation methods, Automatic Segmentation (ASEG) and Sequence Adaptive Multimodal Segmentation (SAMSEG). These are provided by the recently released version 7.1 of the open-source neuroimaging package FreeSurfer, with ASEG being the default segmentation method. First, we use a large sample of participants (n = 1629) distributed across the lifespan (age range = 4-93 years) to assess the within-session test-retest reliability in eight bilateral subcortical structures: amygdala, caudate, hippocampus, lateral ventricles, nucleus accumbens, pallidum, putamen and thalamus. We performed the analyses separately for a sub-sample scanned on a 1.5T Siemens Avanto (n = 774) and a sub-sample scanned on a 3T Siemens Skyra (n = 855). The absolute symmetrized percent differences across the lifespan indicated relatively constant reliability trajectories across age except for the younger children in the Avanto dataset for ASEG. Although both methods showed high reliability (ICC > 0.95), SAMSEG yielded significantly lower volumetric differences between repeated measures for all subcortical segmentations (p < 0.05) and higher spatial overlap in all structures except putamen, which had significantly higher spatial overlap for ASEG. Second, we tested how well each method was able to detect neuroanatomic volumetric change using longitudinal follow up scans (n = 491 for Avanto and n = 245 for Skyra; interscan interval = 1-10 years). Both methods showed excellent ability to detect longitudinal change, but yielded age-trajectories with notable differences for most structures, including the hippocampus and the amygdala. For instance, ASEG hippocampal volumes showed a steady age-decline from subjects in their twenties, while SAMSEG hippocampal volumes were stable until their sixties. Finally, we tested sensitivity of each method to clinically relevant change. We compared annual rate of hippocampal atrophy in a group of cognitively normal older adults (n = 20), patients with mild cognitive impairment (n = 20) and patients with Alzheimer’s disease (n = 20). SAMSEG was more sensitive to detect differences in atrophy between the groups, demonstrating ability to detect clinically relevant longitudinal changes. Both ASEG and SAMSEG were reliable and led to detection of within-person longitudinal change. However, SAMSEG yielded more consistent measurements between repeated scans without a lack of sensitivity to changes.

## 1. Introduction

Automated techniques for whole-brain segmentation have become extremely useful in the study of a range of brain diseases and conditions, such as Alzheimer’s disease (AD) (Chételat, 2018), and also normal conditions such as development (Ostby et al., 2009) and aging (Wonderlick et al., 2009). Automated techniques enable processing of large numbers of magnetic resonance imaging (MRI) scans with limited operator investments, enabling detailed segmentations of brains from large-scale brain imaging initiatives. One of the most extensively used whole-brain segmentation approach is Automatic Segmentation (ASEG) (Fischl et al., 2002), distributed as part of FreeSurfer (http://freesurfer.net/) (Fischl, 2012). FreeSurfer ASEG is a core tool in large-scale neuroimaging projects such as UK Biobank (≈ 40.000 scans to date) (Alfaro-Almagro et al., 2018), ABCD (≈ 10.000 scans to date) (Hagler et al., 2019), ADNI (> 20.000 scans) (Jack et al., 2008), ENIGMA (> 50.000 scans) (Thompson et al., 2020), and Lifebrain (≈ 10.000 scans) (Walhovd et al., 2018). Although the accuracy of automated segmentation techniques such as ASEG is generally high and lead to accurate detection of longitudinal changes (Mulder et al., 2014; Worker et al., 2018), reports have suggested that segmentation accuracy may vary as a function of variables such as age (Wenger et al., 2014) and brain size (Herten et al., 2019; Schoemaker et al., 2016). Hence, continued efforts are undertaken to improve accuracy and reduce bias in the segmentations.

Similar to many other current whole-brain segmentation techniques, ASEG is based on supervised models of T1-weighted scans. As signal intensities alone are not sufficient to distinguish between different neuroanatomical structures from a T1-weighted MRI, an atlas containing probabilistic information about the location of structures is used to determine the relationship between intensities and neuroanatomical labels in particular regions of the brain. The probabilistic atlas is generated from a set of manually labeled training scans. The segmentation problem is then solved in a Bayesian framework in which local shape, position and appearance all contribute to the probability of a given label. Recently, an alternative approach was suggested - Sequence Adaptive Multimodal Segmentation (SAMSEG) – which uses generative parametric models (Puonti et al., 2013, 2016). SAMSEG uses a mesh-based computational atlas combined with a Gaussian appearance model, which is an intensity clustering algorithm that achieves independence of specific image contrast by grouping together voxels with similar intensities (Van Leemput, 2009). SAMSEG is less computationally demanding than other iterative segmentation methods since no preprocessing is needed and only a single, efficient non-linear registration of the atlas to the target image is required. Moreover, bias field estimation and correction is done simultaneous with segmentation and non-linear registration. Nevertheless, SAMSEG resulted in accuracy comparable to ASEG and three other state-of-the-art methods in segmenting T1-weighted MRIs (Puonti et al., 2016). Since SAMSEG does not rely on the specific intensity profiles of a separate training data set, it yields consistent segmentations across scanner platforms and pulse sequences. SAMSEG is included as part of the recent FreeSurfer 7.1 release (released May 11th, 2020), which enables its general use in the neuroimaging community. Therefore, a thorough analysis is necessary to direct many researchers who have the choice between these two utilities provided in the same widely used package of FreeSurfer.

In the present study we undertake a thorough comparative analysis of SAMSEG and ASEG in terms of reliability, bias, sensitivity to longitudinal change, and clinical sensitivity. First, in a large sample of participants (n = 1629) distributed across the lifespan (age range = 4-93 years), we assessed within-session test-retest reliability, including whether this varied with age and structure size. We performed analysis separately for a sub-sample scanned on a 1.5T Siemens Avanto (n = 774) and a sub-sample scanned on a 3T Siemens Skyra (n = 855). Further, since high test-retest reliability could come at the cost of lower sensitivity to biologically meaningful change, we tested how well ASEG and SAMSEG were able to detect neuroanatomic volumetric change in longitudinal follow up scans (n = 491 for Avanto and n = 245 for Skyra; interval between scans = 1-10 years). Finally, we tested how sensitive each method is to clinically relevant change. We compared the annual rate of hippocampal atrophy in a group of cognitively normal older adults (CN) (n = 20), patients with mild cognitive impairment (MCI) (n = 20) and patients with AD (n = 20), and tested the power of each method to detect differences in rates of hippocampal atrophy between the groups.

## 2. Materials and Methods

### 2.1. Datasets

#### 2.1.1. Lifespan scan-rescan dataset

We use scan-rescan data selected from several ongoing projects at the Center for Lifespan Changes in Brain and Cognition (LCBC), University of Oslo, approved by the Regional Committees for Medical and Health Research Ethics South of Norway. Participants were cognitively healthy, and all participants or their guardian provided informed consent (for details, see e.g. (Walhovd et al., 2016)). The images were acquired using two models of Siemens MRI scanners (Siemens Medical Solutions, Erlangen, Germany) - 1.5T Avanto and 3T Skyra, at Rikshospitalet, Oslo University Hospital. A total of 890 participants (1643 sessions) and 887 participants (1739 sessions) were included in the initial within-session scan-rescan datasets for Avanto and Skyra scanners respectively. After discarding scans with insufficient image quality (detailed in section 2.2), the samples were reduced to 774 participants (427 females; 1362 sessions; age range = 4-93 years) for Avanto and 855 participants (563 females; 1646 sessions; age range = 14-84 years) for Skyra. Fig 1 summarizes age distributions for each scanner dataset.

**Fig 1.**
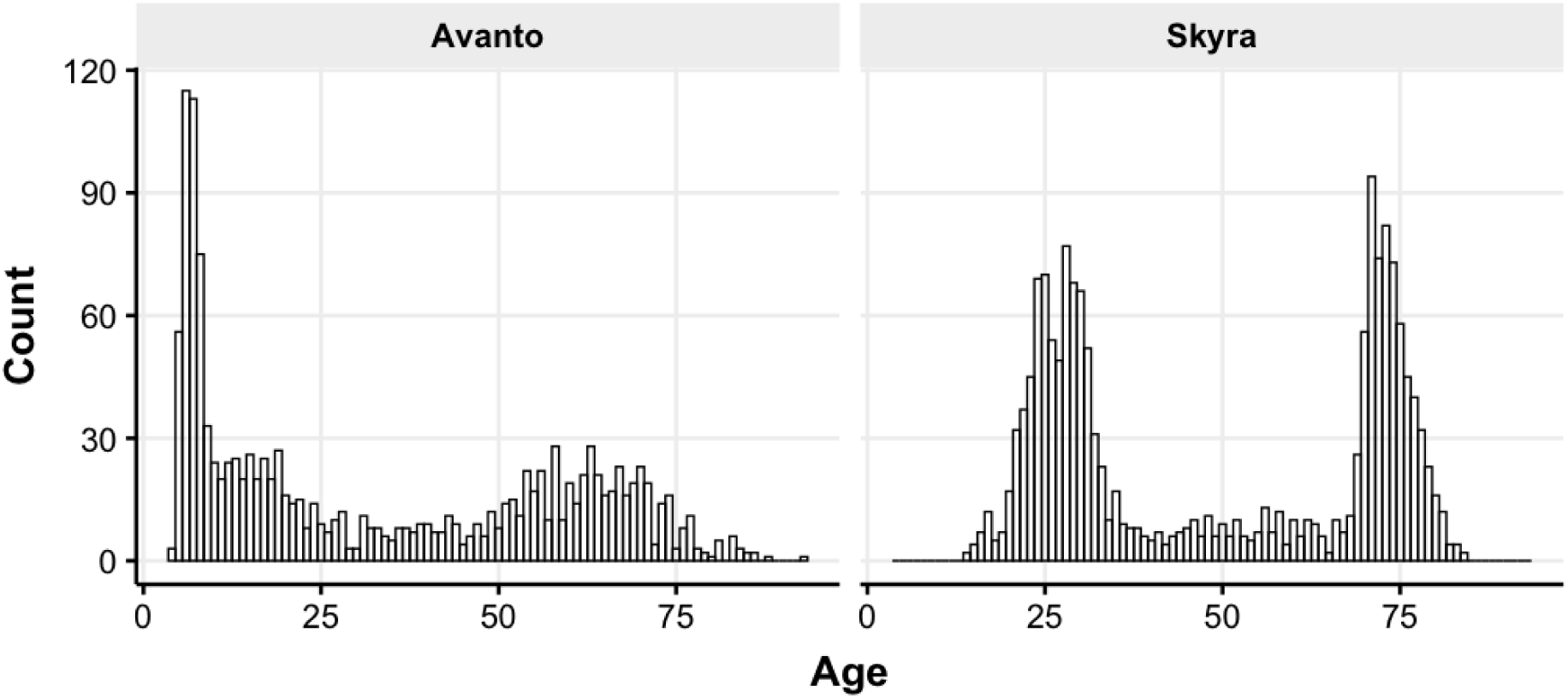
Age distributions of Avanto and Skyra datasets for scan-rescan analysis.

#### 2.1.2. Lifespan longitudinal datasets

For longitudinal LCBC datasets, we selected participants from the scan-rescan dataset who also had a follow-up visit: 491 participants of the Avanto scanner and 245 participants of the Skyra scanner. Each participant had two visits with the follow-up ranging from 1 to 10 years for the Avanto dataset and 1 to 5 years for the Skyra dataset.

#### 2.1.3. Clinical sensitivity dataset

In addition to longitudinal LCBC datasets, we also included scans from the Alzheimer’s disease Neuroimaging Initiative (ADNI) database (adni.loni.usc.edu). The ADNI was launched in 2003 as a public-private partnership, led by Principal Investigator Michael W. Weiner, MD. The primary goal of ANDI has been to test whether serial MRI, positron emission tomography, other biological makers, and clinical and neuropsychological assessment can be combined to measure the progression of MCI and early AD. For up-to-date information, see www.adni-info.org. For our study we randomly selected three groups of participants with similar age distributions: CN, MCI and AD. Each group consisted of 20 participants.

### 2.2. MRI acquisition

MRI acquisition parameters for LCBC samples differed between the scanners but were identical for each session on the same scanner, except for the Skyra dataset where one scan was acquired using parallel acquisition factor GRAPPA = 1 and rescanned with GRAPPA = 2. Avanto data was acquired using Magnetization Prepared Rapid Gradient Echo (MPRAGE) sequence with parameters: TR = 2400 ms, TE = 3.61 ms, TI = 1000 ms, flip angle = 8°, voxel size = 1.25 x 1.25 x 1.2 mm3, 192 x 192 acquisition matrix, 160 slices, 180 Hz pixel bandwidth, GRAPPA = 1, 12 channels head coil. Skyra data was also acquired using MPRAGE sequence with parameters: TR = 2300 ms, TE = 2.98 ms, TI = 850 ms, flip angle = 8°, voxel size = 1.0 x 1.0 x 1.0 mm3, 256 x 256 acquisition matrix, 176 slices, 240 Hz pixel bandwidth, GRAPPA = 1 and 2, 20 channels head coil. Subjects were not repositioned between scan and rescan acquisitions, to acquire data with optimal comparability within each scanner. All images were visually inspected for motion and other artefacts, and sessions that had two scans of acceptable image quality were included in further analysis.

The selected sample of ADNI data was acquired at different sites using a Siemens Avanto 1.5T MRI scanner and MPRAGE sequence: TR = 2400 ms, TE = 3.54 ms, TI = 1000 ms, flip angle = 8°, voxel size = 1.25 x 1.25 x 1.2 mm3, 192 x 192 acquisition matrix, 160 slices, 180 Hz pixel bandwidth, GRAPPA = 1, 8 channel matrix coil. Each participant had two visits with a follow-up ranging from 6 months to 2 years for each group.

### 2.3. MRI processing

Due to the non-linearity of the magnetic fields from the imaging gradient coils, images from each scanner were first preprocessed to reduce geometrical variability of the same subject’s brains between different sessions. This was achieved by obtaining scanner-specific spherical harmonics expansions that represent the gradient coils (Jovicich et al., 2006).

Two fully automated subcortical segmentation methods FreeSurfer v7.1 ASEG and SAMSEG were used to process MRI data and measure volumes of eight bilateral brain structures of interest: amygdala, caudate, hippocampus, lateral ventricles, nucleus accumbens, pallidum, putamen and thalamus. Briefly, the FreeSurfer ASEG pipeline includes Talairach transformation, intensity correction, the removal of nonbrain tissues and volumetric brain segmentation based upon the existence of an atlas containing information on the location of structures, whereas SAMSEG utilizes a mesh-based atlas and a Bayesian modelling framework to obtain volumetric segmentations without the need for skull-stripping as it includes segments for extra cerebral cerebrospinal fluid, skull and soft tissue outside of the skull. Both methods are fully automated and model-based that use a pre-built probabilistic atlas prior from 39 and 20 subjects, respectively. The 20 subjects used for SAMSEG are a subset of the 39 used for ASEG.

To extract reliable volume estimates between scan-rescan acquisitions, we processed images with the longitudinal stream in FreeSurfer ASEG and SAMSEG. For FreeSurfer ASEG, an unbiased within-subject template space and image (Reuter and Fischl, 2011) is created using robust, inverse consistent registration (Reuter et al., 2010). Several processing steps, such as skull stripping, Talairach transforms, atlas registration, and spherical surface maps and parcellations are then initialized with common information from the within-subject template, significantly increasing reliability and statistical power (Reuter et al., 2012). Longitudinal SAMSEG is based on a generative model of longitudinal data (Iglesias et al., 2016). In the forward model, a subject-specific atlas is obtained by generating a random warp from the usual population atlas, and subsequently each time point is again randomly warped from this subject-specific atlas. Bayesian inference is used to obtain the most likely segmentations, with the intermediate subject-specific atlas playing the role of latent variable in the model, whose function is to ensure that various time points have atlas warps that are similar between themselves, without having to define a priori what these warps should be similar to.

### 2.4. Statistical Analysis

#### 2.4.1. Scan-rescan differences

Several statistical approaches were used to describe and evaluate the magnitude of within-session scan-rescan variability between volume measurements. The absolute symmetrized percent difference (ASPD) was calculated:

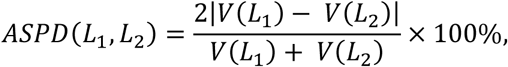

where *V*(*L*) is the volume of the segmented label of structure *L*. ASPD value of 0 indicates a perfect replicability, with increasing values indicating less reliable repeated measurements. Generalized additive models (GAM) (Wood, 2017) were used to characterize volume estimation variability trends of subcortical structures across the lifespan. GAMs are generalized linear models in which the predictors depend linearly or non-linearly on some smooth non-linear functions (Hastie and Tibshirani, 1990). The smooth functions are estimated from the data and enable a flexible smooth curve fitting across the lifespan.

ASPD captures how much the structure differs in terms of its size estimates between repeated measures but does not provide specific insight on the spatial variability. Therefore, a fractional volume overlap or Dice’s coefficient (Dice, 1945) was also computed:

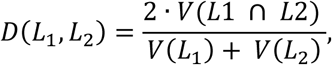

where *V*(*L*1 ∩ *L*2) is the volume of the structure representing the intersection of two labels. In case of two perfectly overlapping structures, Dice coefficient is 1, with decreasing values indicating worse spatial overlap.

Intraclass correlation coefficient (ICC) is a widely used reliability measure for inter-rater, intra-rater and test-retest analyses. It defines the extent to which measurements can be replicated and reflects not only degree of correlation but also agreement between measurements. A value close to 1 indicates a high reliability, with decreasing values indicating lower reliability. ICC estimates and their 95% confidence intervals were calculated using a 2-way mixed-effects model, single measurement and absolute agreement ICC form (McGraw and Wong, 1996; Koo and Li, 2016).

In addition to ICC we used Bland-Altman plots to analyze agreement between two repeated measurements (Bland and Altman, 1986). For each pair of repeated measurements, the x-axis is the mean of both values and y-axis is the percent difference between the two values. Bland-Altman plots facilitate identification of any systematic differences between the measurements regarding the size of the structure.

#### 2.4.2. Sensitivity to longitudinal change

First, to assess whether the estimated lifespan trajectories of the subcortical volumes differed depending on segmentation method, we used General Additive Mixed Models (GAMM) (Wood, 2017). In contrast to GAMs which treat each observation as independent, GAMMs take longitudinal information into account by explicitly modeling the correlation between repeated measurements of the same subject, yielding a model which captures cross-sectional and longitudinal information. Second, to assess longitudinal changes, we used the annualized percentage change (APC) values between the baseline and the follow-up visits for all participants with two scans separated by one year or more. We compared APC values for each segmentation method with paired samples t-tests. We also divided the sample into development (< 20 years), adulthood (between 20 and 60 years) and aging (> 60 years), and compared APCs between each age-group using t-tests and Cohen’s D. Cohen’s D is an effect size used to indicate the standardized difference between two means. Third, to address the clinical sensitivity of each segmentation method, we computed APC for the hippocampus for ADNI subjects, and assessed differences in APC between groups (NC vs. MCI vs. AD) using Cohen’s D. Finally, we used Receiver Operating Characteristic (ROC) curves and Area Under the Curve (AUC) to address the classification sensitivity based on the APC values of the longitudinal hippocampus estimates in different groups.

All statistical analyses described above were done using R statistical software package v3.6.3 (R Core Team, 2020) and its related packages: *mgcv* (Wood, 2017), *ggplot2* (Wickham, 2016), *ggpubr* (Kassambara, 2020), *cowplot* (Wilke, 2019)*, irr* (Gamer et al., 2019), *effsize* (Torchiano, 2020) and *dplyr* (Wickham et al., 2020).

## 3. Results

### 3.1. Scan-rescan reliability

Fig 2 shows volume estimation differences between repeated acquisitions across the lifespan for the Avanto dataset. Although most of the subcortical structures indicated relatively flat lifespan trends, small deviations were observed for ASEG for the young children group which also demonstrated larger variance between the repeated measurements. Fig 3 shows comparable results for the Skyra dataset. However, the Skyra dataset did not include young children and the age-related trends did not show larger differences in the younger sample. Interestingly, the lateral ventricles indicated linearly higher scan-rescan reliability with aging for both scanners and methods. Fig 4 summarizes the overall performance of each segmentation method on both scanner datasets across the lifespan. SAMSEG volume estimates resulted in significantly lower (paired samples t-test, p < 0.05) scan-rescan differences than ASEG for all structures. In addition, the standard deviations were also lower for SAMSEG.

**Fig 2.**
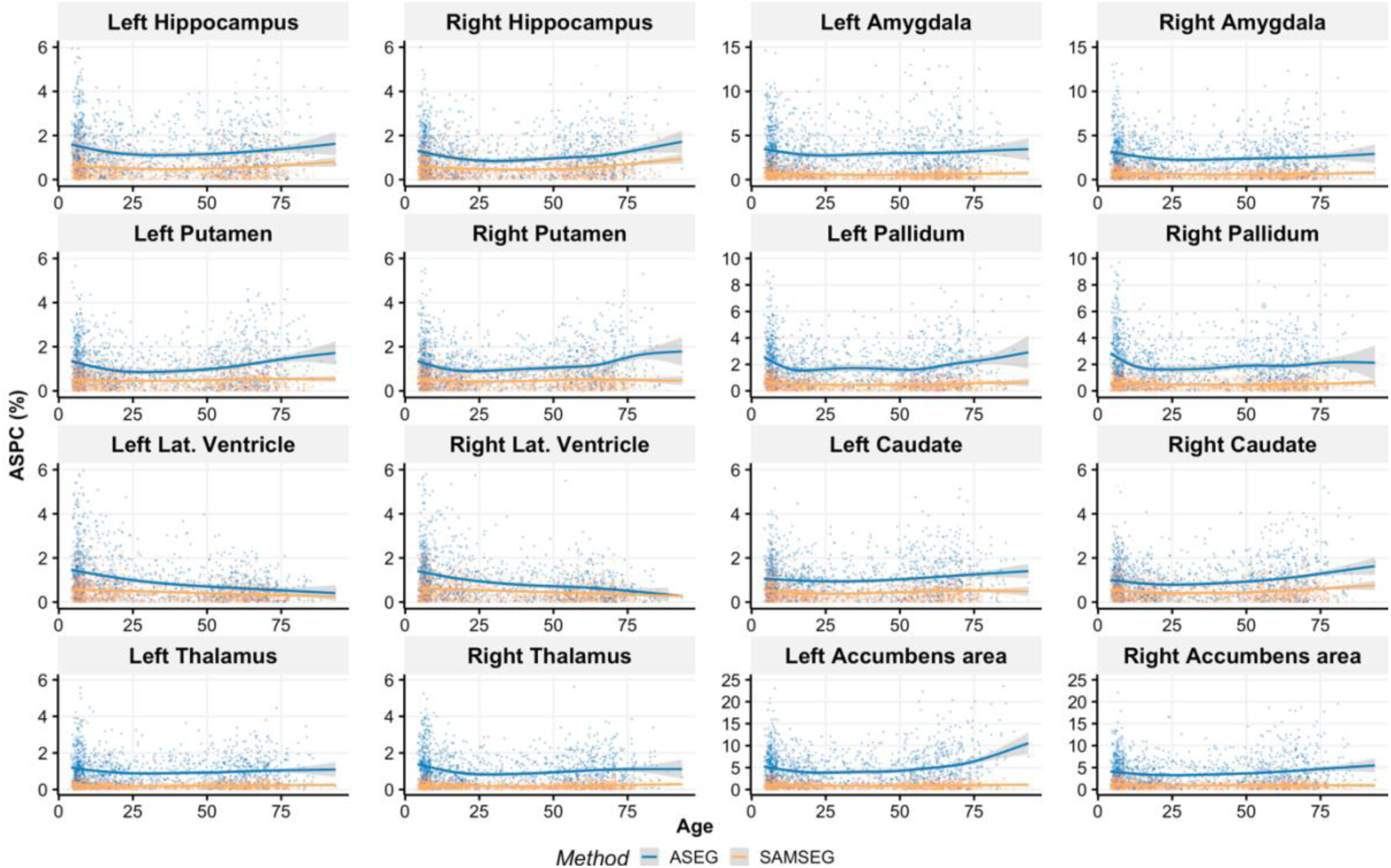
ASPC values across the lifespan for the Avanto dataset. Age-related trends for each method are shown by the GAM curves.

**Fig 3.**
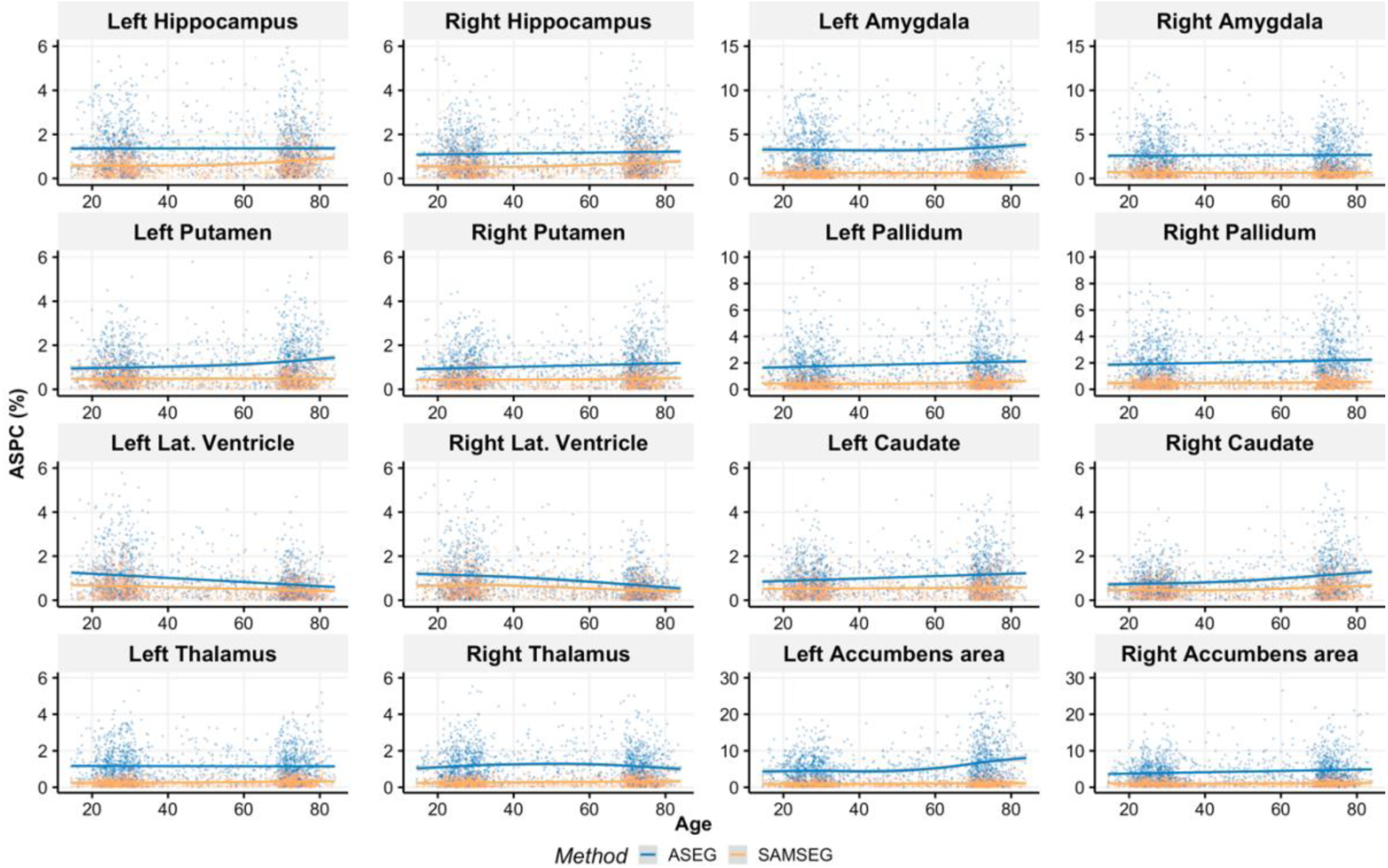
ASPC values across the lifespan for the Skyra dataset. Age-related trends for each method are shown by the GAM curves.

**Fig 4.**
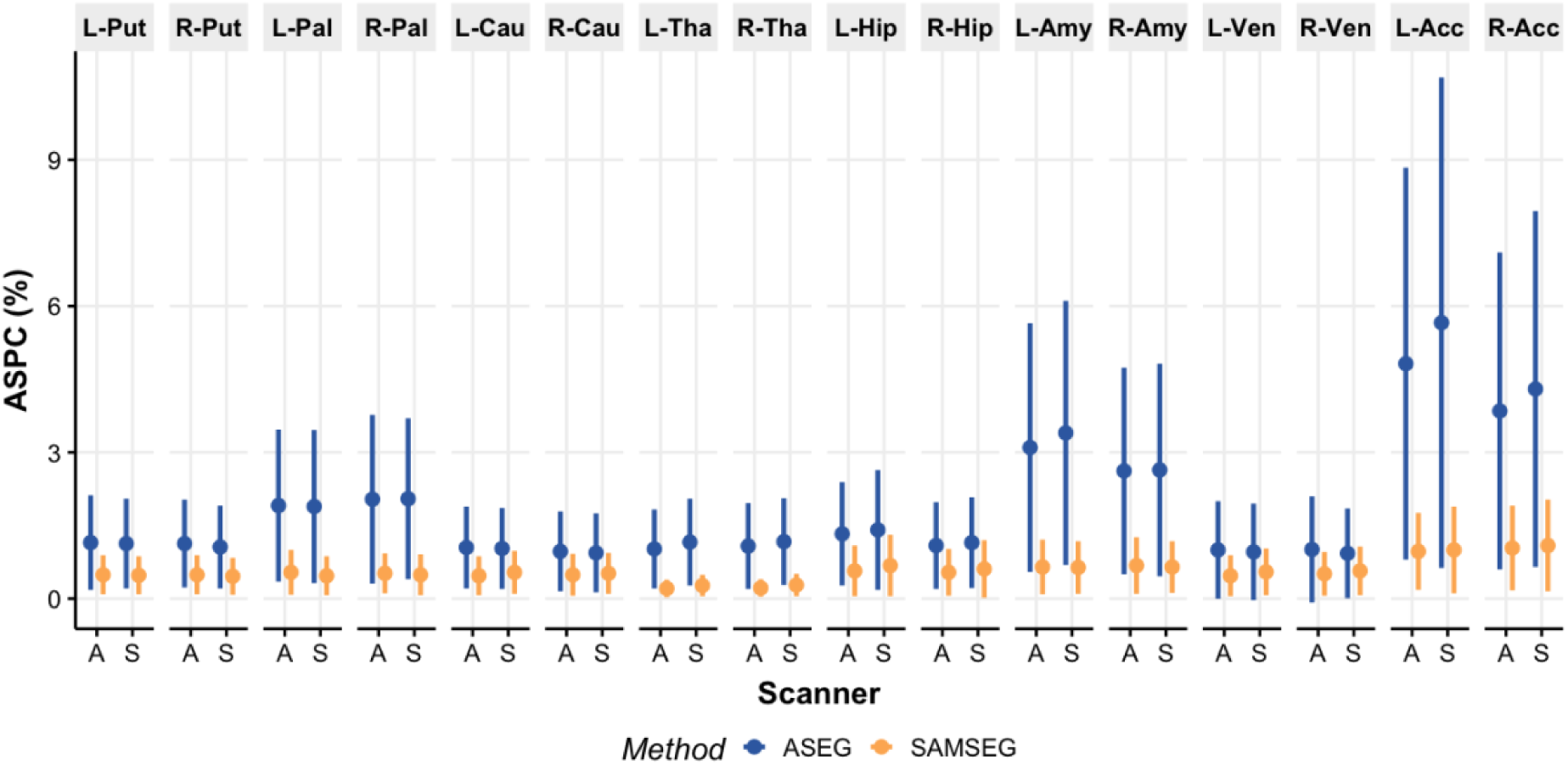
Mean ASPC values (dots) and standard deviations (vertical bars) for each scanner dataset and segmentation method for the subcortical structures: L-Put (left putamen), R-Put (right putamen), L-Pal (left pallidum), R-Pal (right pallidum), L-Cau (left caudate), R-Cau (right caudate), L-Tha (left thalamus), R-Tha (right thalamus), L-Hip (left hippocampus), R-Hip (right hippocampus), L-Amy (left amygdala), R-Amy (right amygdala), L-Ven (left lateral ventricle), R-Ven (right lateral ventricle), L-Acc (left nucleus accumbens), R-Acc (right nucleus accumbens). X-axis indicates Avanto (A) and Skyra (S) scanners.

Fig 5 shows the lifespan test-retest Dice scores for the Avanto dataset. Most of the structures indicated inverted u-shape trajectories except the lateral ventricles which demonstrated almost linearly increasing reliability with aging. Fig 6 illustrates similar Dice scores and age-related trends for the Skyra dataset. However, we found linearly worse spatial overlap with aging because it did not include young children. Fig 7 summarizes the Dice scores across the lifespan. ASEG yielded significantly higher spatial agreement for putamen (both hemispheres and scanners, paired samples t-test, p < 0.01) whereas the rest of the spatial overlaps were significantly better for SAMSEG. The largest improvements were demonstrated for amygdala, pallidum and nucleus accumbens.

**Fig 5.**
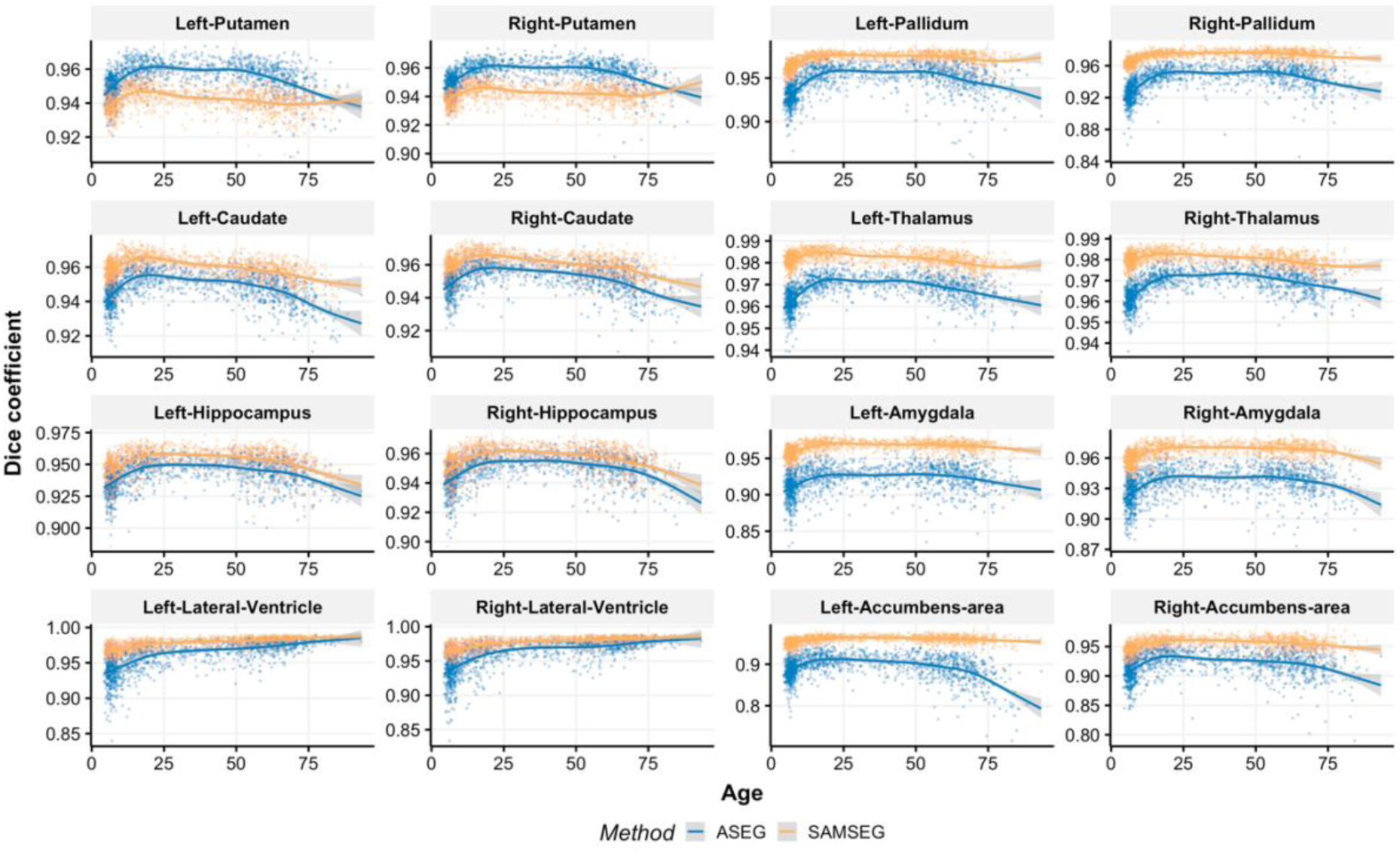
Dice coefficients across the lifespan for the Avanto dataset. Age-related trajectories are shown by the GAM curves. The y-axis scale varies across plots to enable easier evaluation of age-trends.

**Fig 6.**
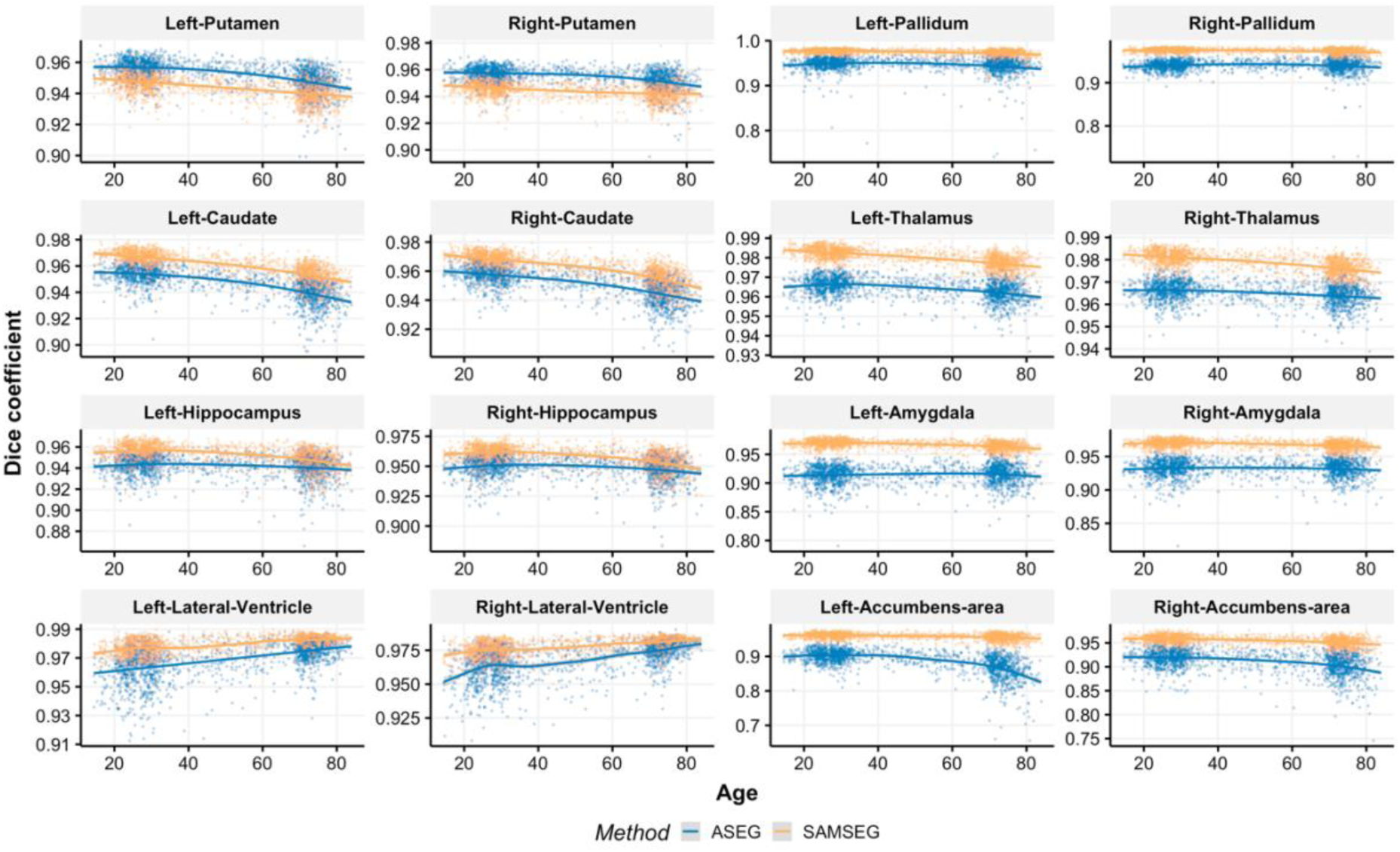
Dice coefficients across the lifespan for the Skyra dataset. Age-related trajectories are shown by the GAM curves. The y-axis scale varies across plots to facilitate easier evaluation of age-trends.

**Fig 7.**
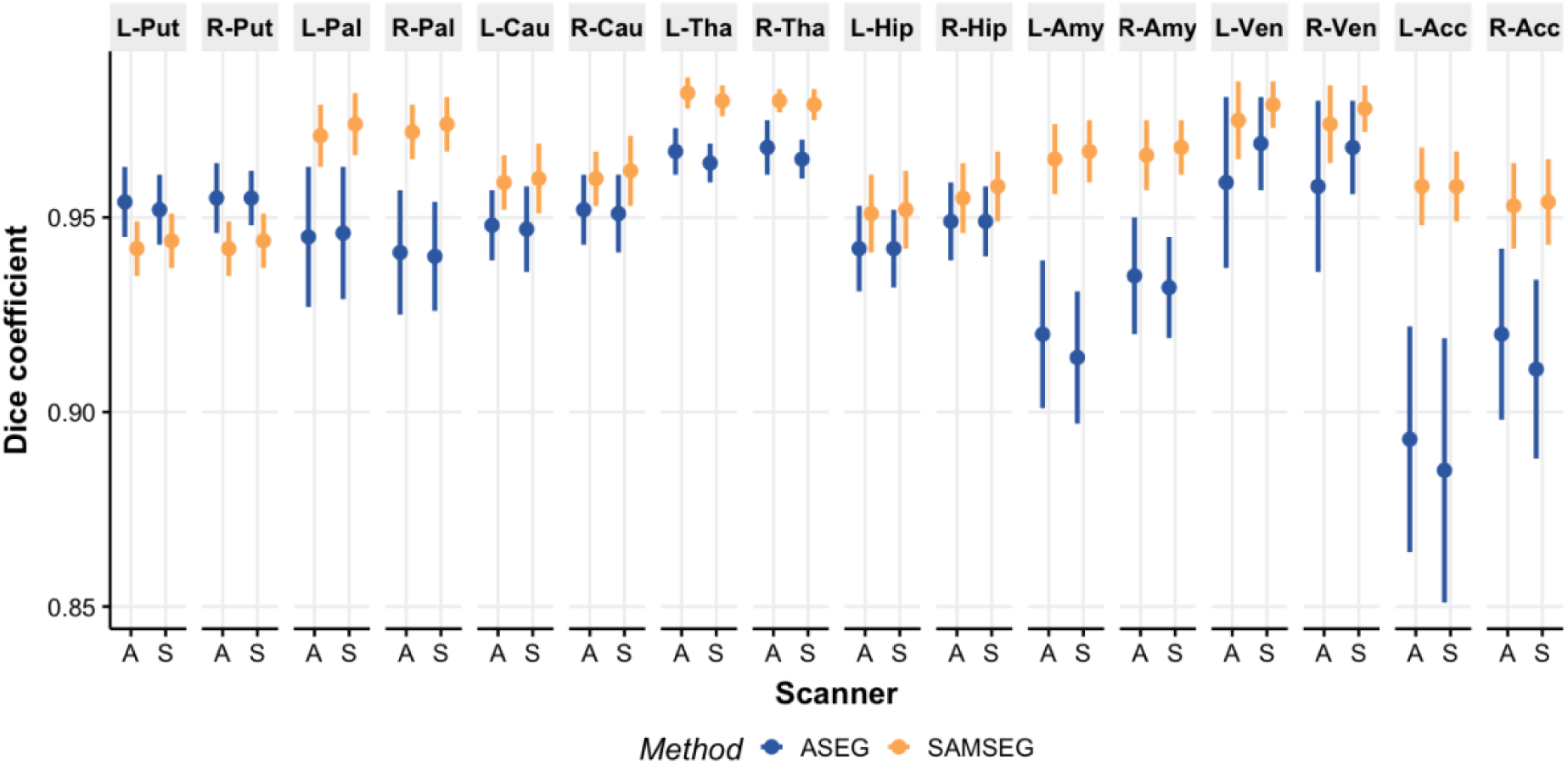
Mean Dice coefficients (dots) and standard deviations (vertical bars) for each scanner dataset and segmentation method for the subcortical structures: L-Put (left putamen), R-Put (right putamen), L-Pal (left pallidum), R-Pal (right pallidum), L-Cau (left caudate), R-Cau (right caudate), L-Tha (left thalamus), R-Tha (right thalamus), L-Hip (left hippocampus), R-Hip (right hippocampus), L-Amy (left amygdala), R-Amy (right amygdala), L-Ven (left lateral ventricle), R-Ven (right lateral ventricle), L-Acc (left nucleus accumbens), R-Acc (right nucleus accumbens). X-axis indicates Avanto (A) and Skyra (S) scanners.

The ICC was computed to assess the agreement between the repeated measurements for each scanner dataset and segmentation method. Although the reliability of the repeated measurements was very high (ICC > 0.95) for both methods, SAMSEG resulted in significantly higher (p < 0.01) ICC values than ASEG for all subcortical structures.

Fig 8 and Fig 9 show Bland-Altman plots for the Avanto dataset. Despite consistent volumetric estimations regardless of the structure size, the limits of agreement *(average difference ± 1.96 standard deviation of the difference)* were in favor of SAMSEG. Smaller lateral ventricles yielded higher variance for both methods. Similar results were observed for the Skyra dataset.

**Fig 8.**
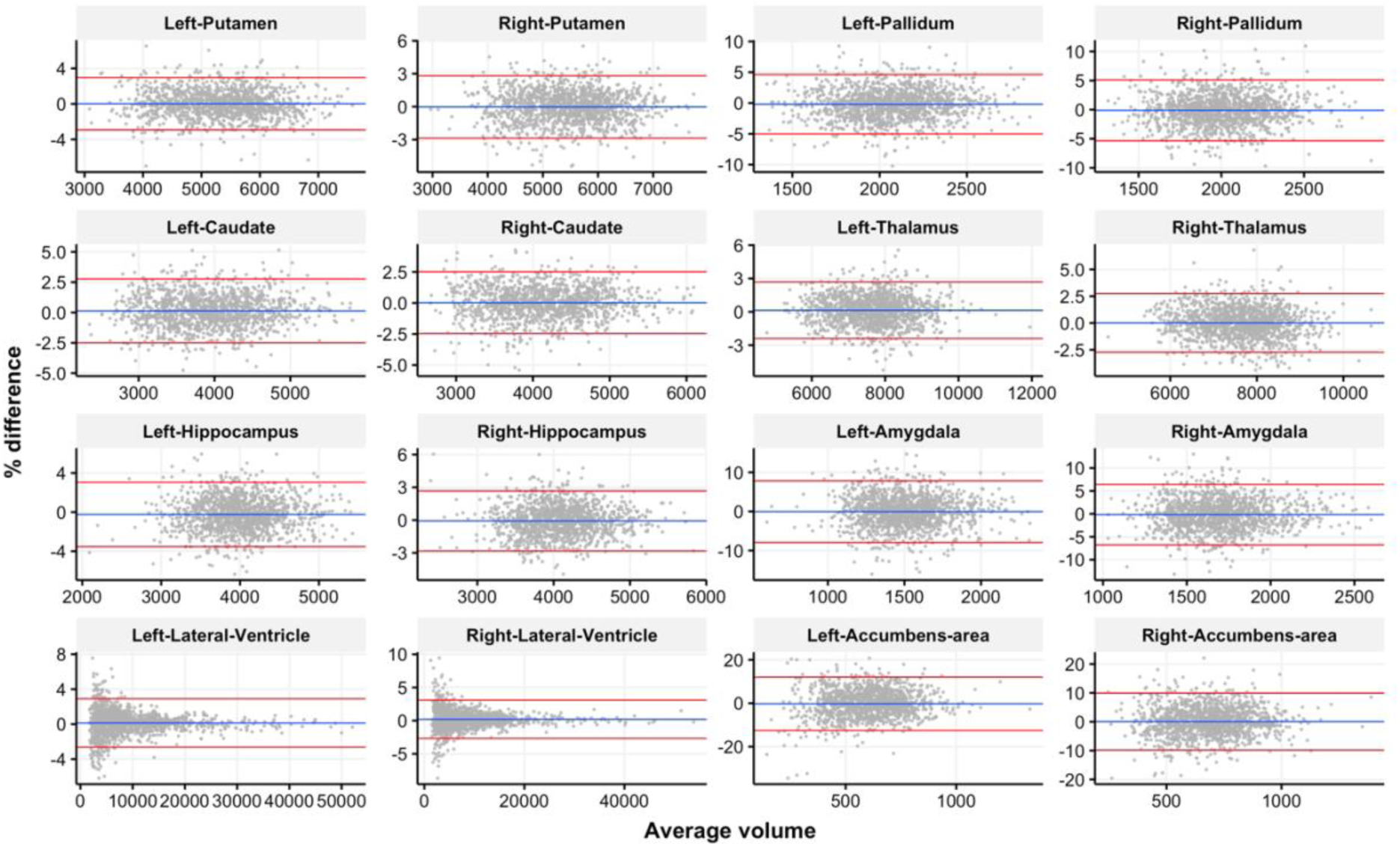
Bland-Altman plots for the Avanto dataset and ASEG segmentation method. Limits of agreement (average difference ± 1.96 standard deviation of the difference) are shown by the red lines.

**Fig 9.**
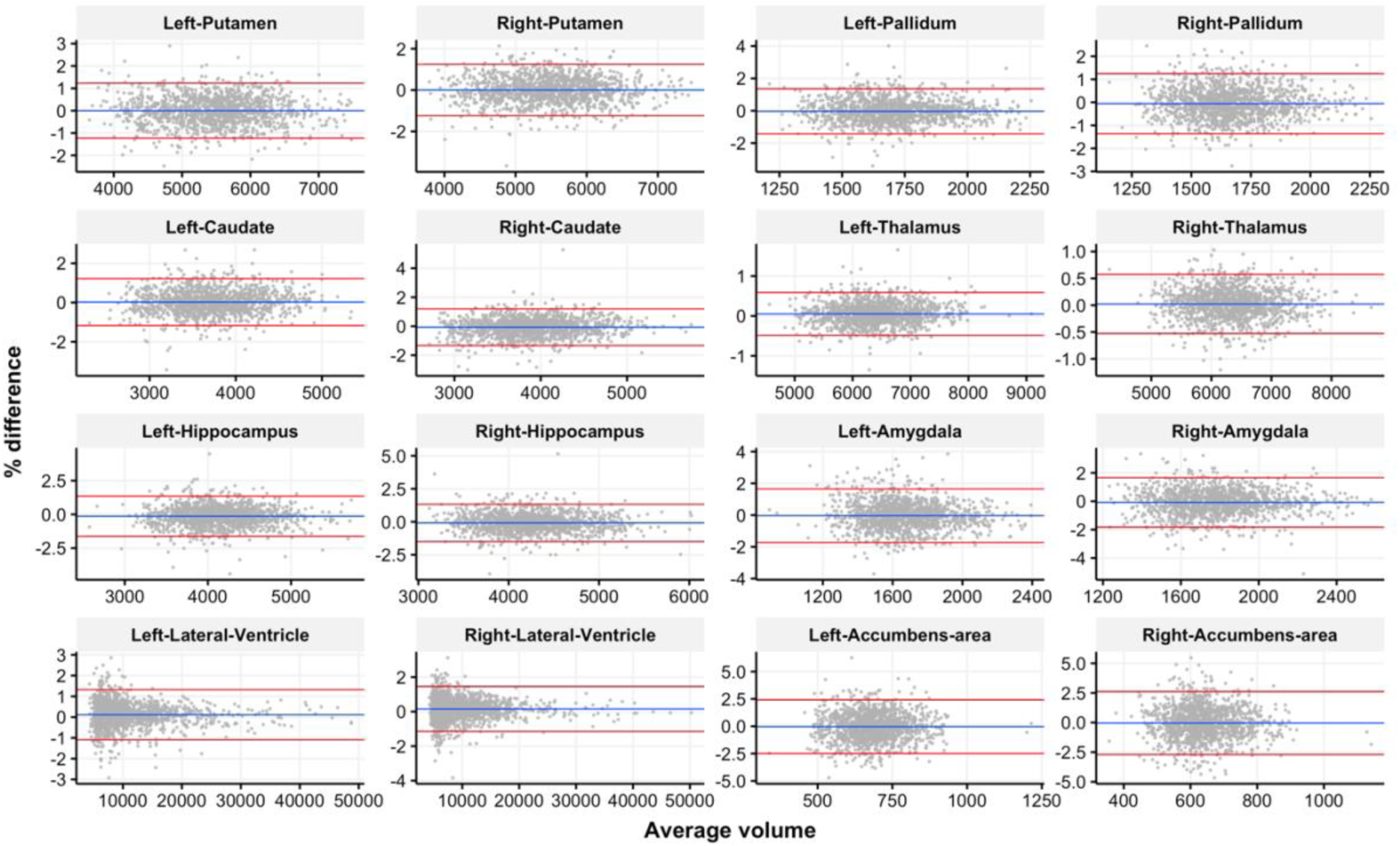
Bland-Altman plots for the Avanto dataset and SAMSEG segmentation method. Limits of agreement (average difference ± 1.96 standard deviation of the difference) are shown by the red lines.

### 3.2. Longitudinal changes

Higher intra-scanner reliability could be a result of lower sensitivity to detect relevant change in brain volumes. We therefore tested the sensitivity of ASEG and SAMSEG to detect changes over time using longitudinal scans and previously documented effects. First, to test whether ASEG and SAMSEG yielded different estimated lifespan trajectories for the volume of each structure when both cross-sectional and longitudinal information was taken into account, we ran GAMMs. For this, we used the part of the LCBC sample where two observations separated by at least one year were available for each participant. Each volume was modelled as a function of age, which would vary within each participant with more than one test occasion. The resulting curves thus take into account both observed within-participant change and between participant differences in age. Fig 10 shows the lifespan trajectories for each method for the Avanto dataset. Although there were similarities in estimated age-trajectories between methods, there were also marked differences. Especially, ASEG estimated more prominent age-effects for the hippocampus, amygdala and thalamus structures, with apparent volumetric reductions starting at a much earlier age compared to the SAMSEG results. Similar observations held for the Skyra dataset.

**Fig 10.**
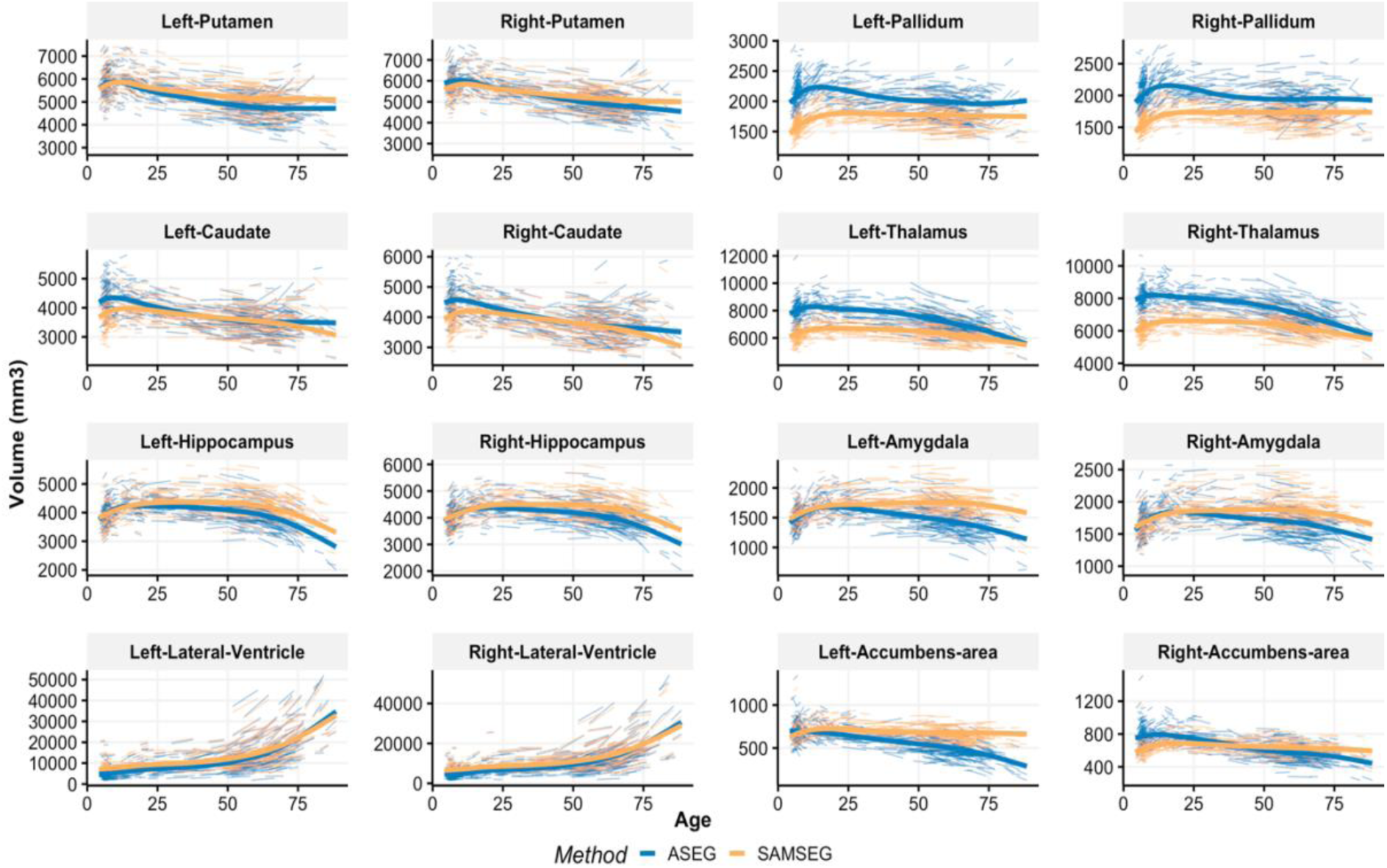
Lifespan trajectories estimated from the combined cross-sectional and longitudinal data for the Avanto scanner and both segmentation methods. Lifespan trajectories are estimated by GAMM, and represent a combination of cross-sectional and longitudinal information.

Next, we analyzed change as indexed by the APC between time-points. The sample was divided into 3 age groups: development, adulthood and aging as described in the Section 2.3. Fig 11 shows the summary of APC values between age groups and segmentation methods for the left and right hippocampus of the Avanto dataset. Hippocampus was chosen because of its known vulnerability both in normal aging and in degenerative diseases such as AD. All estimated mean APC values were significantly different from zero (t-test, p < 0.01) showing that both methods were sensitive to change in all three groups. The standard deviations were also smaller for SAMSEG. Based on paired samples t-tests, the mean differences in the APC values between the segmentation methods for each age group were all significant (p < 0.01) indicating that SAMSEG tended to estimate smaller longitudinal changes than ASEG. Similar results were observed for the Skyra dataset in adulthood and aging groups.

**Fig 11.**
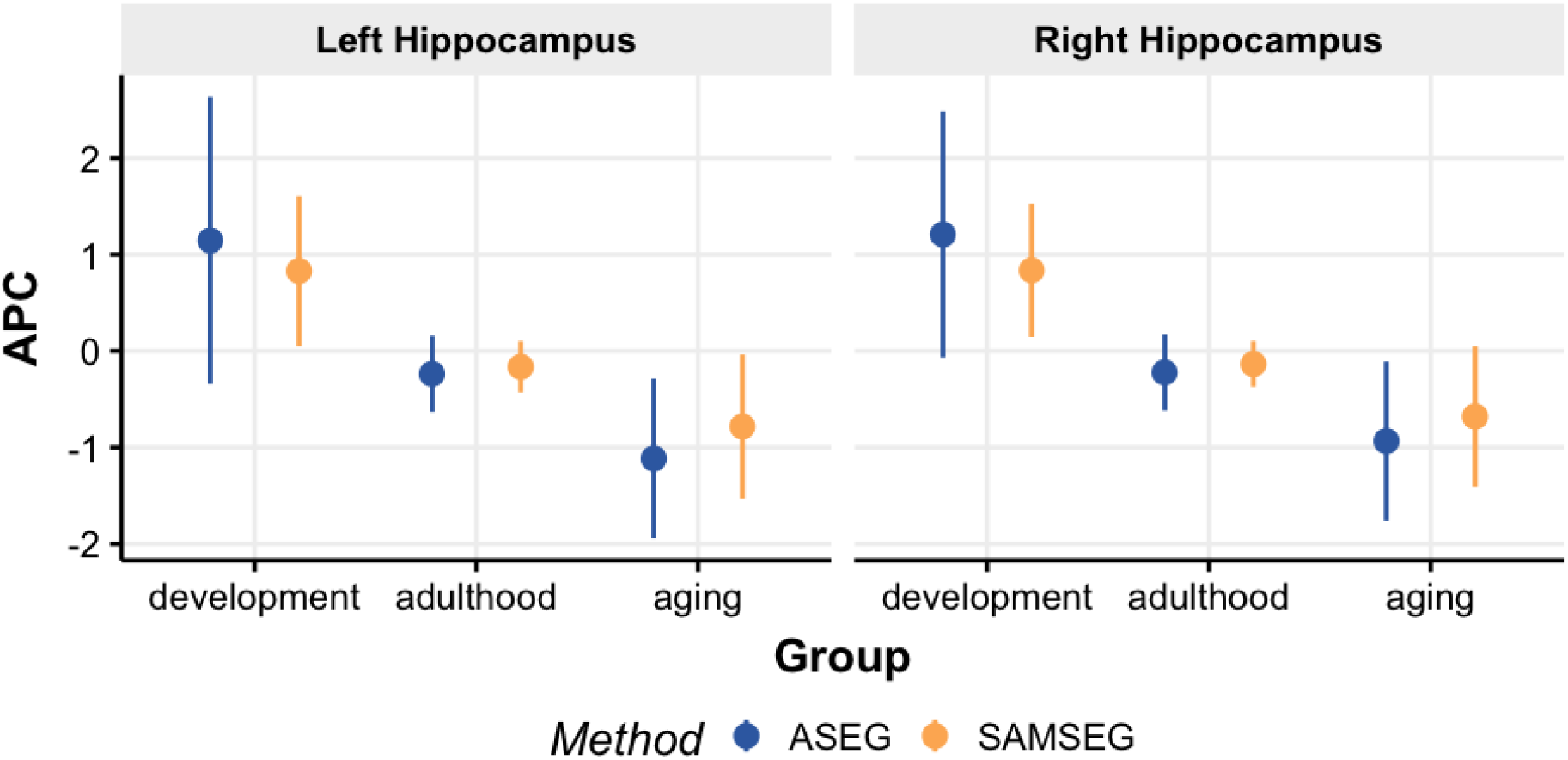
A summary of the mean APC values (dots) and its standard deviations (vertical bars) between age groups for the Avanto dataset.

Fig 12 summarizes the effect sizes (Cohen’s D) based on the APC values between development and adulthood, and between adulthood and aging for the Avanto dataset. SAMSEG yielded larger numeric effect sizes between development and adulthood, and ASEG between adulthood and aging. However, none these differences were significant. Similar results were observed for the Skyra dataset.

**Fig 12.**
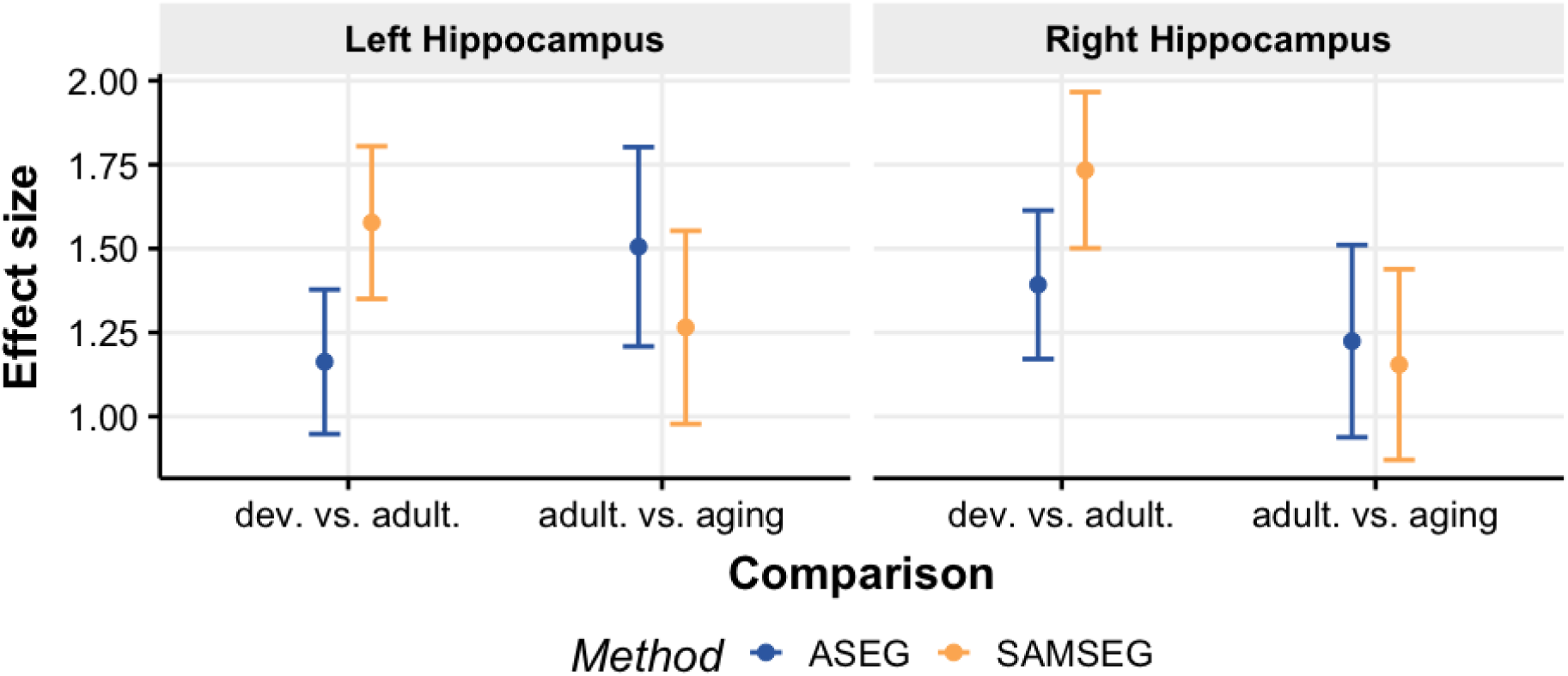
Cohen’s D effect sizes (dots) and their 95% confidence intervals (vertical bars) for development vs. adulthood, and adulthood vs. aging groups for the Avanto dataset.

**Fig 13.**
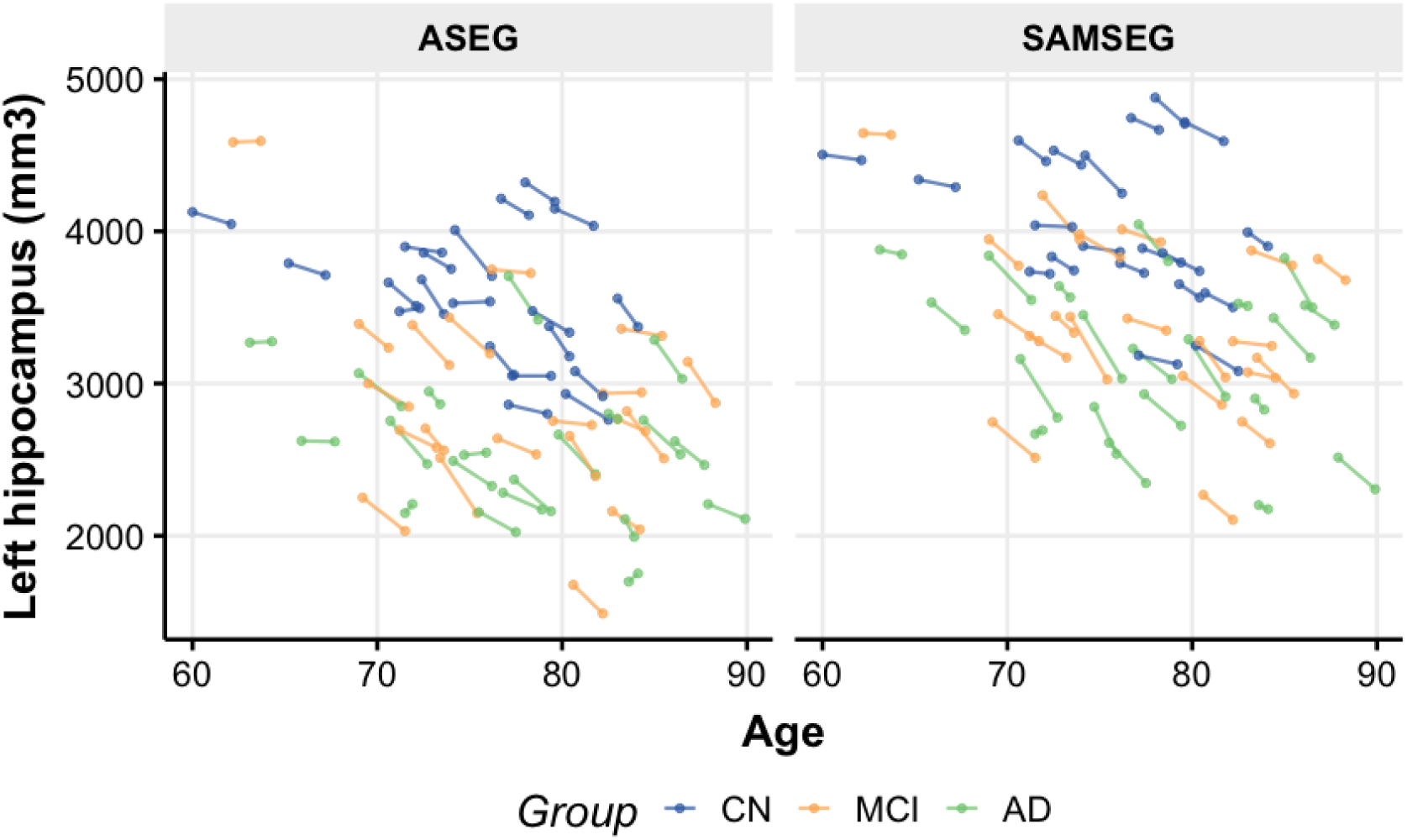
Longitudinal left hippocampus volume changes between the segmentation methods for CN, MCI and AD groups.

### 3.3 Clinical sensitivity

The results of the longitudinal changes indicate that SAMSEG yields lower APC estimates than ASEG, but also smaller standard deviations. However, there is no ground truth whether less or more estimated changes is more accurate. Therefore, we addressed the clinical sensitivity using a subsample of ADNI data. For the purpose of this analysis we only considered a hippocampus since it is the most sensitive structure for detecting AD.

Fig 14 shows longitudinal left hippocampus volume changes. The observed differences were very similar between the methods but SAMSEG yielded larger changes for some of the participants in the AD group. In addition, SAMSEG tended to estimate larger volumes compared to ASEG but this was consistent between the groups.

**Figure 14.**
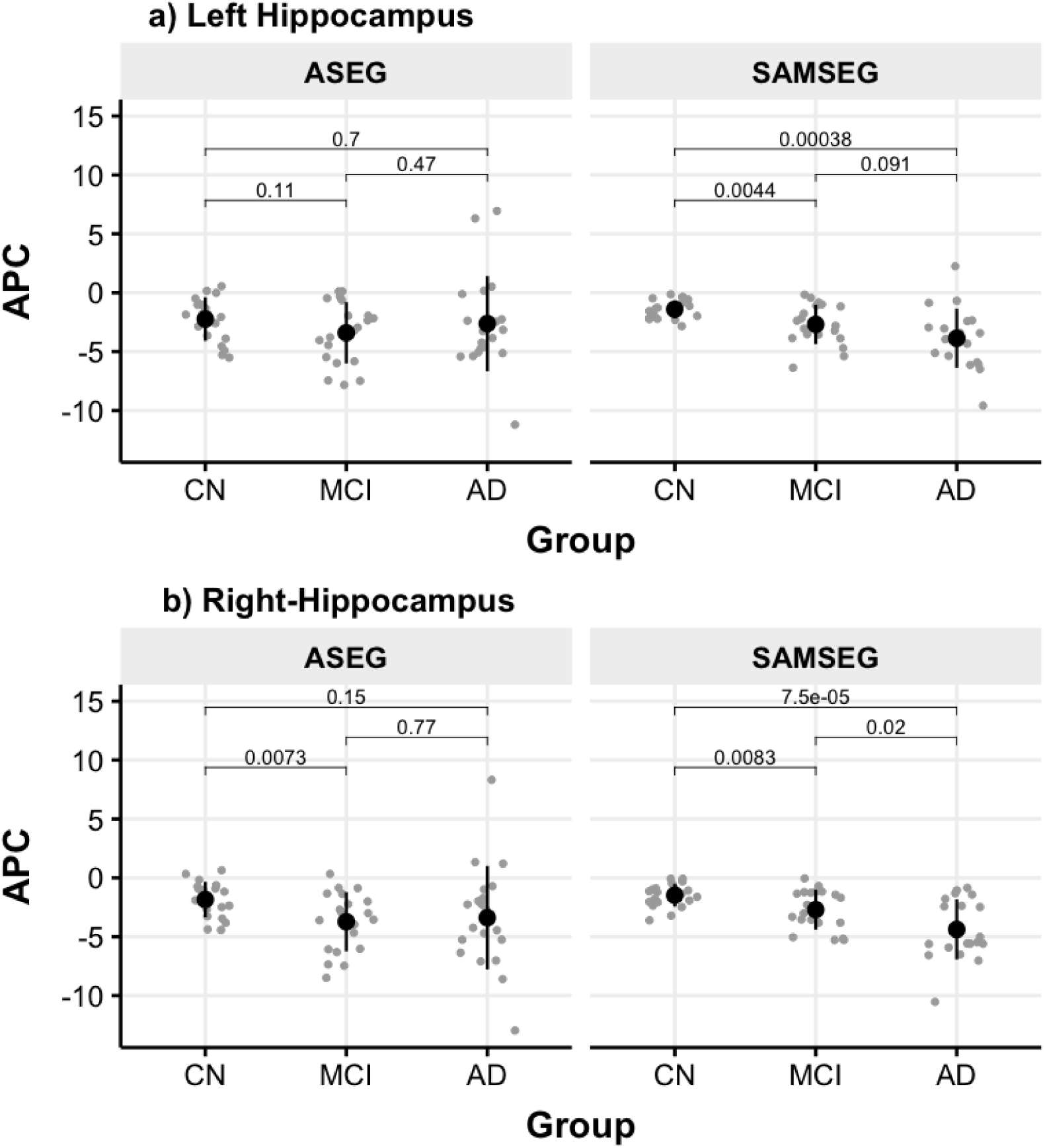
Group comparisons of the estimated a) left and b) right hippocampus APC values from the longitudinal ASEG and SAMSEG segmentations. Group means and standard deviations are shown by the black vertical point range markers. The p-values of the t-tests between the group means are indicated above the horizontal bars.

Fig 14 presents the group comparisons based on the estimated hippocampus APC values from the longitudinal ASEG and SAMSEG segmentations. SAMSEG led to detection of significant differences in atrophy rates between all clinical groups except for the left hippocampus MCI vs. AD comparison. For ASEG, significant differences were seen for the right hippocampus CN vs. MCI contrast. Generally, ASEG demonstrated larger APC variability within each group than SAMSEG.

Fig 15 summarizes the effect sizes (Cohen’s D) and their 95% confidence intervals between the group comparisons. The effects were generally larger for SAMSEG than ASEG but none of these were significantly different between the segmentation methods.

**Fig 15.**
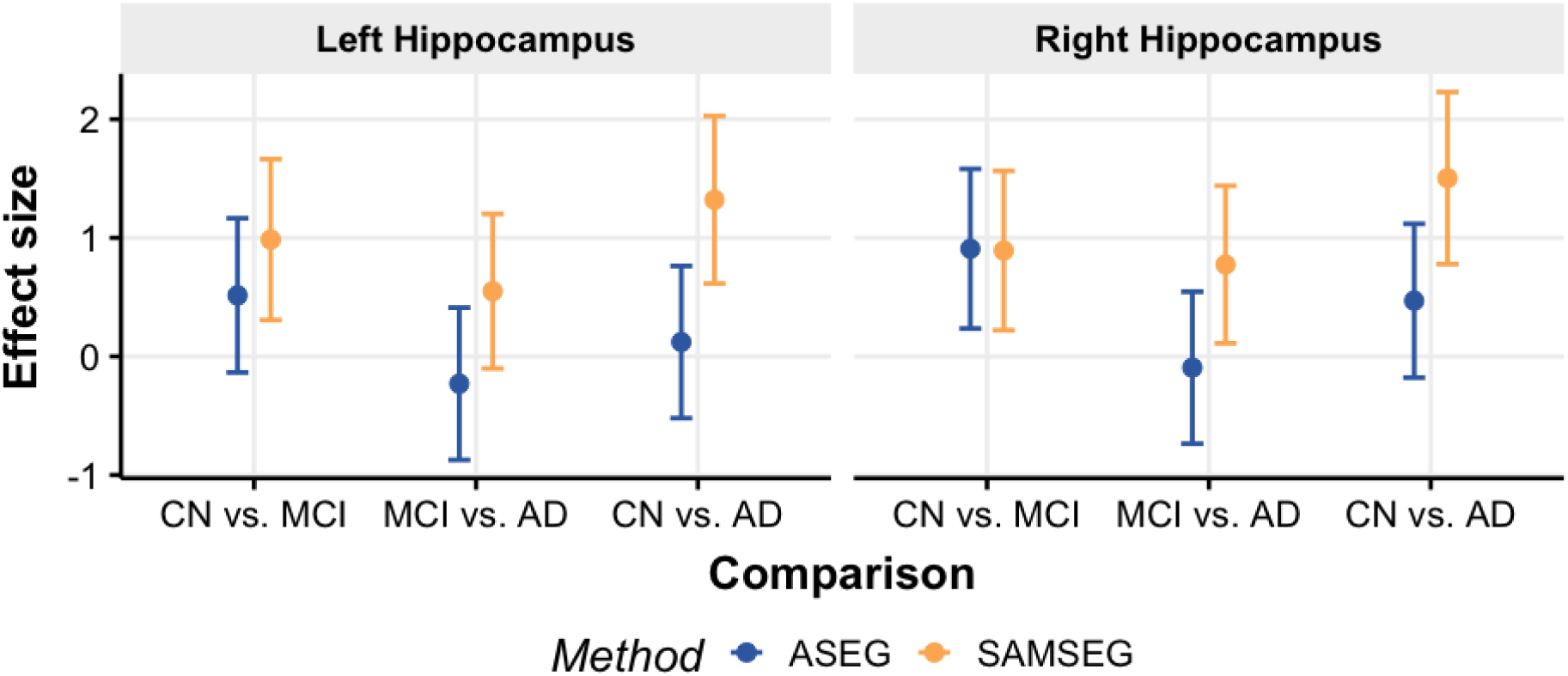
Cohen’s D effect sizes (dots) and their 95% confidence intervals (vertical bars) for the group comparisons between ASEG and SAMSEG for the left and right hippocampus.

Fig 16 plots ROC-AUC curves for classification of patients into different groups based on the APC values of hippocampus. SAMSEG yielded a larger number of correct classifications at the same or lower rate of false positives than ASEG. A very similar scenario was observed for the right hippocampus.

**Fig 16.**
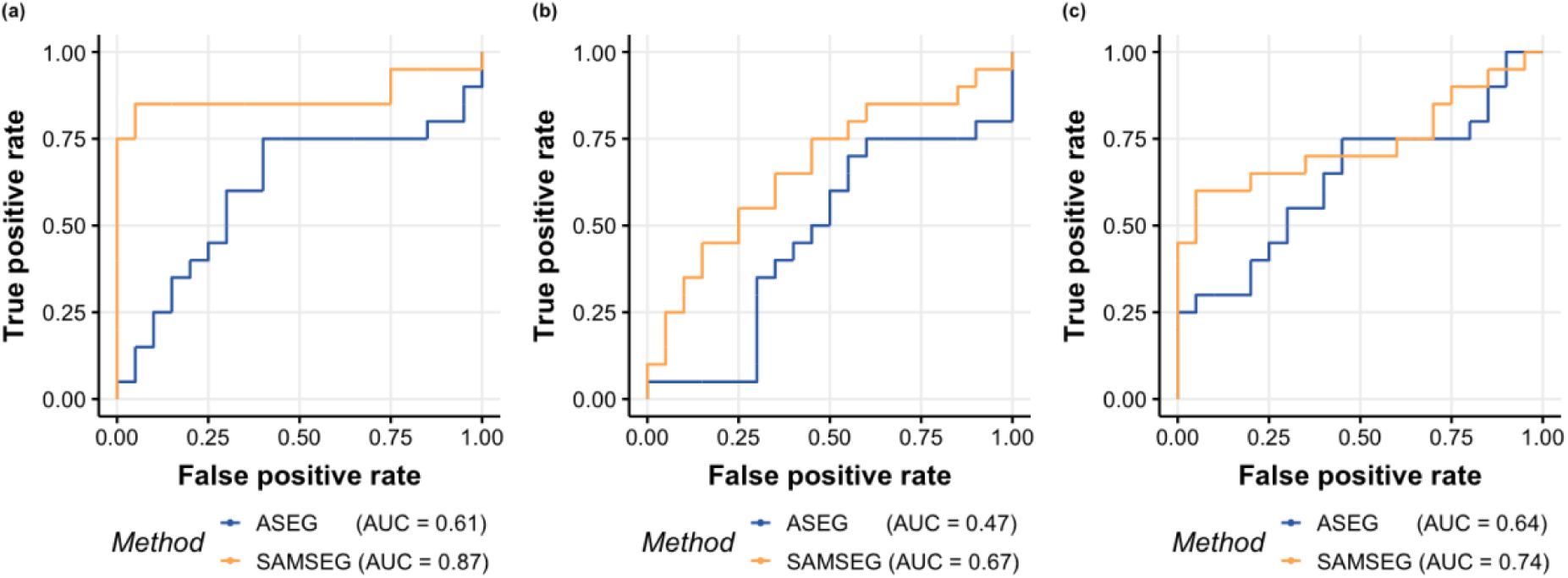
The ROC-AUC curves for classifying participants based on the APC values of the longitudinal hippocampus estimates: (a) AD from CN, (b) AD from MCI and (c) MCI from CN.

## 4. Discussion

The results of the scan-rescan reliability indicated reliable volume estimation across the lifespan, scanners and segmentation methods. Slight deviations were observed for younger participants, presumably due to subtle head motion artifacts. It has previously been shown that younger age groups typically evidence increased motion artifacts, which can hinder the identification of the tissue boundaries (Blumenthal et al., 2002). Importantly, subtle motion artifacts can lead to systematic biases in automatic measurement of structural brain properties (Yendiki et al., 2014). Although different parallel imaging factors (GRAPPA) were used for the Skyra scan-rescan dataset (GRAPPA = 2 vs. GRAPPA = 1), it did not indicate sensitivity to lower signal-to-noise ratio and was comparable to the Avanto dataset. Similar effects of parallel imaging acceleration were shown by (Wonderlick et al., 2009).

The observed average volumetric differences across the lifespan for ASEG were similar to previous reports (Jovicich et al., 2009; Morey et al., 2010). Nevertheless, SAMSEG led to significantly higher scan-rescan volume estimation reliability for all subcortical structures and higher spatial overlap in all structures except putamen, which had significantly higher spatial overlap for ASEG. This is likely a result of SAMSEG’s probabilistic atlas, which currently does not include claustrum structure.

High within-session reliability could come at the cost of less sensitivity to detect meaningful biological change, i.e. that SAMSEG over-regularizes. However, the present analyses of within-person longitudinal change suggest that SAMSEG does ***not*** achieve improved reliability by sacrificing sensitive to change. Both with SAMSEG and with ASEG, longitudinal changes in hippocampal volume were detected, and the APC values were comparable. In the absence of the ground truth longitudinal changes, the present findings suggest that both methods are sensitive to changes in hippocampal volume over time.

We also mapped the lifespan trajectory of each of the structures of interest using GAMMs, taking both cross-sectional and longitudinal information into account. The segmentation differences between ASEG and SAMSEG had substantial effect on lifespan trajectories of all of the tested brain structures, except for the lateral ventricles. In general, developmental trajectories were similar regardless of segmentation method, replicating previous findings (Ostby et al., 2009), although effect sizes for the hippocampus were larger for SAMSEG than ASEG when comparing development to adulthood. For adulthood and aging, however, marked differences were seen for most structures. For the hippocampus and amygdala, the ASEG results replicated earlier studies showing slight volumetric decline from young adulthood (Fjell et al., 2013), with acceleration of volume loss from the sixties, especially marked for the hippocampus. This was not seen for SAMSEG, where very little volume loss was seen before the accelerated decline in aging. For thalamus and pallidum, there were large offset effects, where the estimated volumes for the young children were much higher for ASEG, followed by a steady decline after development ends, extending throughout the rest of the lifespan. This pattern, which is in agreement with previous literature (Fjell et al., 2013), was not seen with SAMSEG. For these structures, as well as nucleus accumbens, SAMSEG yielded modest decline across adulthood, with only some acceleration of volume loss towards the end of life for thalamus. Interestingly, while the previously reported U-shaped trajectory for caudate (Fjell et al., 2013) was seen with ASEG, this was less evident with SAMSEG, which showed a more linear volume decline also in higher age. The implications of these findings await further explorations, but the present results show that the two segmentation methods have substantial effects on the estimated lifespan trajectories of most subcortical structures.

The longitudinal changes analyzed in the clinical setting suggest that SAMSEG tended to be more sensitive to differences in hippocampal atrophy between CN, MCI and AD. This is especially important for detecting the early accelerated hippocampal atrophy which is known to be one of the most sensitive biomarkers of Alzheimer’s disease (Teipel et al., 2013). Expected group differences were more consistently observed for SAMSEG than ASEG. This is likely the result of larger variability between change estimates for ASEG which in turn reduces the power to detect significant differences between the groups. Therefore, based on the current study there is evidence that ASEG might need more samples per group in order to observe the expected group differences, whereas SAMSEG already showed greater sensitivity to detect relevant changes with the relatively modest number of 20 patients in each group that we used for assessment. This is well reflected in the Cohen’s D effect sizes and ROC-AUC curves, which indicate the excellence of classifications based on SAMSEG’s segmentations.

Despite SAMSEG’s high test-retest reliability, it did not indicate reduced sensitivity for biologically meaningful differences. On the contrary, it demonstrated higher sensitivity to detect longitudinal changes than ASEG between development and adulthood, and in the clinical setting.

We analyzed scan-rescan reliability of participants that were not repositioned before acquiring a repeated scan. This scenario is unlikely in the clinical setting where the participants are usually taken out of the scanner before acquiring another repeated scan. This, in turn, might lead to an increased measurement variability and less reliable volumetric estimates compared to what was observed in the present work. Finally, we performed a comprehensive evaluation of longitudinal changes and sensitivity for the hippocampus structure. The remaining subcortical structures should be addressed in addition as it is not evident that similar longitudinal trends would be present.

## 5. Conclusions

Both whole-brain segmentation methods ASEG and SAMSEG demonstrated high test-retest reliability and did not indicate bias towards age (except young children) or structure size. Nevertheless, the reliability measures of SAMSEG were significantly higher for all subcortical structures. Although SAMSEG yielded more consistent measurements between repeated scans, this did not indicate a lack of sensitivity to detect changes. On the contrary, both ASEG and SAMSEG led to detection of within-person longitudinal change, while we found greater sensitivity to detect longitudinal and clinically relevant changes with SAMSEG compared to ASEG. Therefore, the method demonstrates a potential widespread application of the new whole-brain segmentation in the neuroimaging research community. The present findings will also direct many researchers who have the choice between these two utilities, leading to a downstream impact in clinical studies and laying the foundation for further studies that can build on this.

## 6. Acknowledgement

The present research was funded by a grant from Helse-Sør Øst (grant number 2018009) and A.M.F., the European Research Council under grant agreements 283634, 725025 (to A.M.F.), 313440 (to K.B.W.) and 677697 (to J.E.I.), as well as the Norwegian Research Council (to A.M.F., K.B.W.). Data collection and sharing for this project was also funded by the Alzheimer’s Disease Neuroimaging Initiative (ADNI) (National Institutes of Health Grant U01 AG024904) and DOD ADNI (Department of Defense award number W81XWH-12-2-0012). ADNI is funded by the National Institute on Aging, the National Institute of Biomedical Imaging and Bioengineering, and through generous contributions from the following: AbbVie, Alzheimer’s Association; Alzheimer’s Drug Discovery Foundation; Araclon Biotech; BioClinica, Inc.; Biogen; Bristol-Myers Squibb Company; CereSpir, Inc.; Cogstate; Eisai Inc.; Elan Pharmaceuticals, Inc.; Eli Lilly and Company; EuroImmun; F. Hoffmann-La Roche Ltd and its affiliated company Genentech, Inc.; Fujirebio; GE Healthcare; IXICO Ltd.; Janssen Alzheimer Immunotherapy Research & Development, LLC.; Johnson & Johnson Pharmaceutical Research & Development LLC.; Lumosity; Lundbeck; Merck & Co., Inc.; Meso Scale Diagnostics, LLC.; NeuroRx Research; Neurotrack Technologies; Novartis Pharmaceuticals Corporation; Pfizer Inc.; Piramal Imaging; Servier; Takeda Pharmaceutical Company; and Transition Therapeutics. The Canadian Institutes of Health Research is providing funds to support ADNI clinical sites in Canada. Private sector contributions are facilitated by the Foundation for the National Institutes of Health (www.fnih.org). The grantee organization is the Northern California Institute for Research and Education, and the study is coordinated by the Alzheimer’s Therapeutic Research Institute at the University of Southern California. ADNI data are disseminated by the Laboratory for Neuro Imaging at the University of Southern California. The study was also supported by the National Institutes of Health under grant numbers U24DA041123, R01AG057672, R01NR010827 and R01AG059011 (to D.N.G.).

Support for this research was provided in part by the BRAIN Initiative Cell Census Network grant U01MH117023, the National Institute for Biomedical Imaging and Bioengineering (P41EB015896, 1R01EB023281, R01EB006758, R21EB018907, R01EB019956), the National Institute on Aging (1R56AG064027, 1R01AG064027, 5R01AG008122, R01AG016495), the National Institute of Mental Health the National Institute of Diabetes and Digestive and Kidney Diseases (1-R21-DK-108277-01), the National Institute for Neurological Disorders and Stroke (R01NS0525851, R21NS072652, R01NS070963, R01NS083534, 5U01NS086625,5U24NS10059103, R01NS105820, R01NS112161), and was made possible by the resources provided by Shared Instrumentation Grants 1S10RR023401, 1S10RR019307, and 1S10RR023043. Additional support was provided by the NIH Blueprint for Neuroscience Research (5U01-MH093765), part of the multi-institutional Human Connectome Project. In addition, BF has a financial interest in CorticoMetrics, a company whose medical pursuits focus on brain imaging and measurement technologies. BF’s interests were reviewed and are managed by Massachusetts General Hospital and Partners HealthCare in accordance with their conflict of interest policies.

## References

Alfaro-Almagro, F., Jenkinson, M., Bangerter, N.K., Andersson, J.L.R., Griffanti, L., Douaud, G., Sotiropoulos, S.N., Jbabdi, S., Hernandez-Fernandez, M., Vallee, E., Vidaurre, D., Webster, M., McCarthy, P., Rorden, C., Daducci, A., Alexander, D.C., Zhang, H., Dragonu, I., Matthews, P.M., Miller, K.L., Smith, S.M., 2018. Image processing and Quality Control for the first 10,000 brain imaging datasets from UK Biobank. NeuroImage 166, 400–424. https://doi.org/10.1016/j.neuroimage.2017.10.034

Bland, J.M., Altman, D.G., 1986. Statistical methods for assessing agreement between two methods of clinical measurement. Lancet 1, 307–310.

Blumenthal, J.D., Zijdenbos, A., Molloy, E., Giedd, J.N., 2002. Motion Artifact in Magnetic Resonance Imaging: Implications for Automated Analysis. NeuroImage 16, 89–92. https://doi.org/10.1006/nimg.2002.1076

Chételat, G., 2018. Multimodal Neuroimaging in Alzheimer’s Disease: Early Diagnosis, Physiopathological Mechanisms, and Impact of Lifestyle. JAD 64, S199–S211. https://doi.org/10.3233/JAD-179920

Dice, L.R., 1945. Measures of the Amount of Ecologic Association Between Species. Ecology 26, 297–302. https://doi.org/10.2307/1932409

Fischl, B., 2012. FreeSurfer. NeuroImage 62, 774–781. https://doi.org/10.1016/j.neuroimage.2012.01.021

Fischl, B., Salat, D.H., Busa, E., Albert, M., Dieterich, M., Haselgrove, C., van der Kouwe, A., Killiany, R., Kennedy, D., Klaveness, S., Montillo, A., Makris, N., Rosen, B., Dale, A.M., 2002. Whole brain segmentation: automated labeling of neuroanatomical structures in the human brain. Neuron 33, 341–355. https://doi.org/10.1016/s0896-6273(02)00569-x

Fjell, A.M., Westlye, L.T., Grydeland, H., Amlien, I., Espeseth, T., Reinvang, I., Raz, N., Holland, D., Dale, A.M., Walhovd, K.B., 2013. Critical ages in the life course of the adult brain: nonlinear subcortical aging. Neurobiology of Aging 34, 2239–2247. https://doi.org/10.1016/j.neurobiolaging.2013.04.006

Gamer, M., Lemon, J., Singh, I.F.P., 2019. irr: Various Coefficients of Interrater Reliability and Agreement.

Hagler, D.J., Hatton, SeanN., Cornejo, M.D., Makowski, C., Fair, D.A., Dick, A.S., Sutherland, M.T., Casey, B.J., Barch, D.M., Harms, M.P., Watts, R., Bjork, J.M., Garavan, H.P., Hilmer, L., Pung, C.J., Sicat, C.S., Kuperman, J., Bartsch, H., Xue, F., Heitzeg, M.M., Laird, A.R., Trinh, T.T., Gonzalez, R., Tapert, S.F., Riedel, M.C., Squeglia, L.M., Hyde, L.W., Rosenberg, M.D., Earl, E.A., Howlett, K.D., Baker, F.C., Soules, M., Diaz, J., de Leon, O.R., Thompson, W.K., Neale, M.C., Herting, M., Sowell, E.R., Alvarez, R.P., Hawes, S.W., Sanchez, M., Bodurka, J., Breslin, F.J., Morris, A.S., Paulus, M.P., Simmons, W.K., Polimeni, J.R., van der Kouwe, A., Nencka, A.S., Gray, K.M., Pierpaoli, C., Matochik, J.A., Noronha, A., Aklin, W.M., Conway, K., Glantz, M., Hoffman, E., Little, R., Lopez, M., Pariyadath, V., Weiss, S.RB., Wolff-Hughes, D.L.,, DelCarmen-Wiggins, R., Feldstein Ewing, S.W., Miranda-Dominguez, O., Nagel, B.J., Perrone, A.J., Sturgeon, D.T., Goldstone, A., Pfefferbaum, A., Pohl, K.M., Prouty, D., Uban, K., Bookheimer, S.Y., Dapretto, M., Galvan, A., Bagot, K., Giedd, J., Infante, M.A., Jacobus, J., Patrick, K., Shilling, P.D., Desikan, R., Li, Y., Sugrue, L., Banich, M.T., Friedman, N., Hewitt, J.K., Hopfer, C., Sakai, J., Tanabe, J., Cottler, L.B., Nixon, S.J., Chang, L., Cloak, C., Ernst, T., Reeves, G., Kennedy, D.N., Heeringa, S., Peltier, S., Schulenberg, J., Sripada, C., Zucker, R.A., Iacono, W.G., Luciana, M., Calabro, F.J., Clark, D.B., Lewis, D.A., Luna, B., Schirda, C., Brima, T., Foxe, J.J., Freedman, E.G., Mruzek, D.W., Mason, M.J., Huber, R., McGlade, E., Prescot, A., Renshaw, P.F., Yurgelun-Todd, D.A.,, Allgaier, N.A., Dumas, J.A., Ivanova, M., Potter, A., Florsheim, P., Larson, C., Lisdahl, K., Charness, M.E., Fuemmeler, B., Hettema, J.M., Maes, H.H., Steinberg, J., Anokhin, A.P., Glaser, P., Heath, A.C., Madden, P.A., Baskin-Sommers, A., Constable, R.T., Grant, S.J., Dowling, G.J., Brown, S.A., Jernigan, T.L., Dale, A.M., 2019. Image processing and analysis methods for the Adolescent Brain Cognitive Development Study. NeuroImage 202, 116091. https://doi.org/10.1016/j.neuroimage.2019.116091

Hastie, T.J., Tibshirani, R.J., 1990. Generalized Additive Models. CRC Press.

Herten, A., Konrad, K., Krinzinger, H., Seitz, J., von Polier, G.G., 2019. Accuracy and bias of automatic hippocampal segmentation in children and adolescents. Brain Struct Funct 224, 795–810. https://doi.org/10.1007/s00429-018-1802-2

Iglesias, J.E., Van Leemput, K., Augustinack, J., Insausti, R., Fischl, B., Reuter, M., 2016. Bayesian longitudinal segmentation of hippocampal substructures in brain MRI using subject-specific atlases. NeuroImage 141, 542–555. https://doi.org/10.1016/j.neuroimage.2016.07.020

Jack, C.R., Bernstein, M.A., Fox, N.C., Thompson, P., Alexander, G., Harvey, D., Borowski, B., Britson, P.J., L. Whitwell, J., Ward, C., Dale, A.M., Felmlee, J.P., Gunter, J.L., Hill, D.L.G., Killiany, R., Schuff, N., Fox-Bosetti, S., Lin, C., Studholme, C., DeCarli, C.S., Gunnar Krueger, Ward, H.A., Metzger, G.J., Scott, K.T., Mallozzi, R., Blezek, D., Levy, J., Debbins, J.P., Fleisher, A.S., Albert, M., Green, R., Bartzokis, G., Glover, G., Mugler, J., Weiner, M.W., ADNI Study, 2008. The Alzheimer’s disease neuroimaging initiative (ADNI): MRI methods. J. Magn. Reson. Imaging 27, 685–691. https://doi.org/10.1002/jmri.21049

Jovicich, J., Czanner, S., Greve, D., Haley, E., van der Kouwe, A., Gollub, R., Kennedy, D., Schmitt, F., Brown, G., MacFall, J., Fischl, B., Dale, A., 2006. Reliability in multi-site structural MRI studies: Effects of gradient non-linearity correction on phantom and human data. NeuroImage 30, 436–443. https://doi.org/10.1016/j.neuroimage.2005.09.046

Jovicich, J., Czanner, S., Han, X., Salat, D., van der Kouwe, A., Quinn, B., Pacheco, J., Albert, M., Killiany, R., Blacker, D., 2009. MRI-derived measurements of human subcortical, ventricular and intracranial brain volumes: Reliability effects of scan sessions, acquisition sequences, data analyses, scanner upgrade, scanner vendors and field strengths. NeuroImage 46, 177–192. https://doi.org/10.1016/j.neuroimage.2009.02.010

Kassambara, A., 2020. ggpubr: “ggplot2” Based Publication Ready Plots.

Koo, T.K., Li, M.Y., 2016. A Guideline of Selecting and Reporting Intraclass Correlation Coefficients for Reliability Research. J Chiropr Med 15, 155–163. https://doi.org/10.1016/j.jcm.2016.02.012

McGraw, K.O., Wong, S.P., 1996. Forming inferences about some intraclass correlation coefficients. Psychological Methods 1, 30–46. https://doi.org/10.1037/1082-989X.1.1.30

Morey, R.A., Selgrade, E.S., Wagner, H.R., Huettel, S.A., Wang, L., McCarthy, G., 2010. Scan-rescan reliability of subcortical brain volumes derived from automated segmentation. Hum. Brain Mapp. NA-NA. https://doi.org/10.1002/hbm.20973

Mulder, E.R., de Jong, R.A., Knol, D.L., van Schijndel, R.A., Cover, K.S., Visser, P.J., Barkhof, F., Vrenken, H., 2014. Hippocampal volume change measurement: Quantitative assessment of the reproducibility of expert manual outlining and the automated methods FreeSurfer and FIRST. NeuroImage 92, 169–181. https://doi.org/10.1016/j.neuroimage.2014.01.058

Ostby, Y., Tamnes, C.K., Fjell, A.M., Westlye, L.T., Due-Tonnessen, P., Walhovd, K.B., 2009. Heterogeneity in Subcortical Brain Development: A Structural Magnetic Resonance Imaging Study of Brain Maturation from 8 to 30 Years. Journal of Neuroscience 29, 11772–11782. https://doi.org/10.1523/JNEUROSCI.1242-09.2009

Puonti, O., Iglesias, J.E., Van Leemput, K., 2016. Fast and sequence-adaptive whole-brain segmentation using parametric Bayesian modeling. NeuroImage 143, 235–249. https://doi.org/10.1016/j.neuroimage.2016.09.011

Puonti, O., Iglesias, J.E., Van Leemput, K., 2013. Fast, Sequence Adaptive Parcellation of Brain MR Using Parametric Models, in: Salinesi, C., Norrie, M.C., Pastor, Ó. (Eds.), Advanced Information Systems Engineering, Lecture Notes in Computer Science. Springer Berlin Heidelberg, Berlin, Heidelberg, pp. 727–734. https://doi.org/10.1007/978-3-642-40811-3_91

R Core Team, 2020. R: A Language and Environment for Statistical Computing. R Foundation for Statistical Computing, Vienna, Austria.

Reuter, M., Fischl, B., 2011. Avoiding asymmetry-induced bias in longitudinal image processing. NeuroImage 57, 19–21. https://doi.org/10.1016/j.neuroimage.2011.02.076

Reuter, M., Rosas, H.D., Fischl, B., 2010. Highly accurate inverse consistent registration: A robust approach. NeuroImage 53, 1181–1196. https://doi.org/10.1016/j.neuroimage.2010.07.020

Reuter, M., Schmansky, N.J., Rosas, H.D., Fischl, B., 2012. Within-subject template estimation for unbiased longitudinal image analysis. NeuroImage 61, 1402–1418. https://doi.org/10.1016/j.neuroimage.2012.02.084

Schoemaker, D., Buss, C., Head, K., Sandman, C.A., Davis, E.P., Chakravarty, M.M., Gauthier, S., Pruessner, J.C., 2016. Hippocampus and amygdala volumes from magnetic resonance images in children: Assessing accuracy of FreeSurfer and FSL against manual segmentation. NeuroImage 129, 1–14. https://doi.org/10.1016/j.neuroimage.2016.01.038

Teipel, S.J., Grothe, M., Lista, S., Toschi, N., Garaci, F.G., Hampel, H., 2013. Relevance of Magnetic Resonance Imaging for Early Detection and Diagnosis of Alzheimer Disease. Medical Clinics of North America, Early Diagnosis and Intervention in Predementia Alzheimer’s Disease 97, 399–424. https://doi.org/10.1016/j.mcna.2012.12.013

Thompson, P.M., Jahanshad, N., Ching, C.R.K., Salminen, L.E., Thomopoulos, S.I., Bright, J., Baune, B.T., Bertolín, S., Bralten, J., Bruin, W.B., Bülow, R., Chen, J., Chye, Y., Dannlowski, U., de Kovel, C.G.F., Donohoe, G., Eyler, L.T., Faraone, S.V., Favre, P., Filippi, C.A., Frodl, T., Garijo, D., Gil, Y., Grabe, H.J., Grasby, K.L., Hajek, T., Han, L.K.M., Hatton, S.N., Hilbert, K., Ho, T.C., Holleran, L., Homuth, G., Hosten, N., Houenou, J., Ivanov, I., Jia, T., Kelly, S., Klein, M., Kwon, J.S., Laansma, M.A., Leerssen, J., Lueken, U., Nunes, A., Neill, J.O., Opel, N., Piras, Fabrizio, Piras, Federica, Postema, M.C., Pozzi, E., Shatokhina, N., Soriano-Mas, C., Spalletta, G., Sun, D., Teumer, A., Tilot, A.K., Tozzi, L., van der Merwe, C., Van Someren, E.J.W., van Wingen, G.A., Völzke, H., Walton, E., Wang, L., Winkler, A.M., Wittfeld, K., Wright, M.J., Yun, J.-Y., Zhang, G., Zhang-James, Y., Adhikari, B.M., Agartz, I., Aghajani, M., Aleman, A., Althoff, R.R., Altmann, A., Andreassen, O.A., Baron, D.A., Bartnik-Olson, B.L.,, Marie Bas-Hoogendam, J., Baskin-Sommers, A.R.,, Bearden, C.E., Berner, L.A., Boedhoe, P.S.W., Brouwer, R.M., Buitelaar, J.K., Caeyenberghs, K., Cecil, C.A.M., Cohen, R.A., Cole, J.H., Conrod, P.J., De Brito, S.A., de Zwarte, S.M.C., Dennis, E.L., Desrivieres, S., Dima, D., Ehrlich, S., Esopenko, C., Fairchild, G., Fisher, S.E., Fouche, J.-P., Francks, C., Frangou, S., Franke, B., Garavan, H.P., Glahn, D.C., Groenewold, N.A., Gurholt, T.P., Gutman, B.A., Hahn, T., Harding, I.H., Hernaus, D., Hibar, D.P., Hillary, F.G., Hoogman, M., Hulshoff Pol, H.E., Jalbrzikowski, M., Karkashadze, G.A., Klapwijk, E.T., Knickmeyer, R.C., Kochunov, P., Koerte, I.K., Kong, X.-Z., Liew, S.-L., Lin, A.P., Logue, M.W., Luders, E., Macciardi, F., Mackey, S., Mayer, A.R., McDonald, C.R., McMahon, A.B., Medland, S.E., Modinos, G., Morey, R.A., Mueller, S.C., Mukherjee, P., Namazova-Baranova, L., Nir, T.M., Olsen, A., Paschou, P., Pine, D.S., Pizzagalli, F., Rentería, M.E., Rohrer, J.D., Sämann, P.G., Schmaal, L., Schumann, G., Shiroishi, M.S., Sisodiya, S.M., Smit, D.J.A., Sønderby, I.E., Stein, D.J., Stein, J.L., Tahmasian, M., Tate, D.F., Turner, J.A., van den Heuvel, O.A., van der Wee, N.J.A., van der Werf, Y.D., van Erp, T.G.M., van Haren, N.E.M., van Rooij, D., van Velzen, L.S., Veer, I.M., Veltman, D.J., Villalon-Reina, J.E.,, Walter, H., Whelan, C.D., Wilde, E.A., Zarei, M., Zelman, V., 2020. ENIGMA and global neuroscience: A decade of large-scale studies of the brain in health and disease across more than 40 countries. Translational Psychiatry 10, 1–28. https://doi.org/10.1038/s41398-020-0705-1

Torchiano, M., 2020. effsize: Efficient Effect Size Computation.

Van Leemput, K., 2009. Encoding Probabilistic Brain Atlases Using Bayesian Inference. IEEE Trans. Med. Imaging 28, 822–837. https://doi.org/10.1109/TMI.2008.2010434

Walhovd, K.B., Fjell, A.M., Westerhausen, R., Nyberg, L., Ebmeier, K.P., Lindenberger, U., Bartrés-Faz, D., Baaré, W.F.C., Siebner, H.R., Henson, R., Drevon, C.A., Strømstad Knudsen, G.P., Ljøsne, I.B., Penninx, B.W.J.H., Ghisletta, P., Rogeberg, O., Tyler, L., Bertram, L., Lifebrain Consortium, 2018. Healthy minds 00-100 years: Optimising the use of European brain imaging cohorts (“Lifebrain”). Eur. psychiatr. 50, 47–56. https://doi.org/10.1016/j.eurpsy.2017.12.006

Walhovd, K.B., Krogsrud, S.K., Amlien, I.K., Bartsch, H., Bjørnerud, A., Due-Tønnessen, P., Grydeland, H., Hagler, D.J., Håberg, A.K., Kremen, W.S., Ferschmann, L., Nyberg, L., Panizzon, M.S., Rohani, D.A., Skranes, J., Storsve, A.B., Sølsnes, A.E., Tamnes, C.K., Thompson, W.K., Reuter, C., Dale, A.M., Fjell, A.M., 2016. Neurodevelopmental origins of lifespan changes in brain and cognition. Proc Natl Acad Sci USA 113, 9357–9362. https://doi.org/10.1073/pnas.1524259113

Wenger, E., Mårtensson, J., Noack, H., Bodammer, N.C., Kühn, S., Schaefer, S., Heinze, H.-J., Düzel, E., Bäckman, L., Lindenberger, U., Lövdén, M., 2014. Comparing manual and automatic segmentation of hippocampal volumes: Reliability and validity issues in younger and older brains. Human Brain Mapping 35, 4236–4248. https://doi.org/10.1002/hbm.22473

Wickham, H., 2016. ggplot2: Elegant Graphics for Data Analysis. Springer-Verlag New York.

Wickham, H., François, R., Henry, L., Müller, K., 2020. dplyr: A Grammar of Data Manipulation.

Wilke, C.O., 2019. cowplot: Streamlined Plot Theme and Plot Annotations for “ggplot2.”

Wonderlick, J., Ziegler, D., Hosseinivarnamkhasti, P., Locascio, J., Bakkour, A., Vanderkouwe, A., Triantafyllou, C., Corkin, S., Dickerson, B., 2009. Reliability of MRI-derived cortical and subcortical morphometric measures: Effects of pulse sequence, voxel geometry, and parallel imaging. NeuroImage 44, 1324–1333. https://doi.org/10.1016/j.neuroimage.2008.10.037

Wood, S.N., 2017. Generalized Additive Models: An Introduction with R, 2nd ed. Chapman and Hall/CRC.

Worker, A., Dima, D., Combes, A., Crum, W.R., Streffer, J., Einstein, S., Mehta, M.A., Barker, G.J., Williams, S.C.R., O’daly, O., 2018. Test-retest reliability and longitudinal analysis of automated hippocampal subregion volumes in healthy ageing and Alzheimer’s disease populations. Hum Brain Mapp 39, 1743–1754. https://doi.org/10.1002/hbm.23948

Yendiki, A., Koldewyn, K., Kakunoori, S., Kanwisher, N., Fischl, B., 2014. Spurious group differences due to head motion in a diffusion MRI study. NeuroImage 88, 79–90. https://doi.org/10.1016/j.neuroimage.2013.11.027

